# Cactus: a user-friendly and reproducible ATAC-Seq and mRNA-Seq analysis pipeline for data preprocessing, differential analysis, and enrichment analysis

**DOI:** 10.1101/2023.05.11.540110

**Authors:** Jérôme Salignon, Lluís Millan-Ariño, Maxime Garcia, Christian G. Riedel

## Abstract

The ever decreasing cost of Next-Generation Sequencing coupled with the emergence of efficient and reproducible analysis pipelines has rendered genomic methods more accessible. However, downstream analyses are basic or missing in most workflows, creating a significant barrier for non-bioinformaticians. To help close this gap, we developed Cactus, an end-to-end pipeline for analyzing ATAC-Seq and mRNA-Seq data, either separately or jointly. Its Nextflow-, container-, and virtual environment-based architecture ensures efficient and reproducible analyses. Cactus preprocesses raw reads, conducts differential analyses between conditions, and performs enrichment analyses in various databases, including DNA-binding motifs, ChIP-Seq binding sites, chromatin states, and ontologies. We demonstrate the utility of Cactus in a multi-modal and multi-species case study as well as by showcasing its unique capabilities as compared to other ATAC-Seq pipelines. In conclusion, Cactus can assist researchers in gaining comprehensive insights from chromatin accessibility and gene expression data in a quick, user-friendly, and reproducible manner.

## Introduction

The decreasing cost of next-generation sequencing has resulted in omics-scale assays slowly becoming as affordable as conventional assays that only query a few molecules or loci [1]. However, the high dimensionality of omics data remains a challenge. To address this issue, analysis pipelines are useful tools as they reduce the time, expertise, and effort needed by laboratories to analyze their omics-scale data. Specifically, modern pipelines often provide a high degree of reproducibility and efficiency [2] through the use of workflow engines such as Nextflow [3] or SnakeMake [4] and package managers, such as Docker [5], Singularity [6], or conda [7].

mRNA-Seq [8] and ATAC-Seq [9] are two commonly used omics assays because they are affordable, do not require large amounts of sample material, and provide a broad overview of the transcriptome and the regulatory landscape of a sample. mRNA-Seq measures gene expression, while ATAC-Seq measures chromatin accessibility, both without the need to optimize the assay for a specific set of molecules or genomic regions. Many pipelines exist to analyze mRNA-Seq and ATAC-Seq data, but most of them only go as far as peak calling for ATAC-Seq [10–12] or differential gene expression analysis for mRNA-Seq [13,14], leaving many researchers in need of deeper biological insights. Particularly, combining omics results from different methods and performing enrichment analyses is often done in a low-throughput, time-consuming, and hard-to-reproduce manner, creating a strong need for pipelines that offer easy automation of these steps.

To address this, we present Cactus (<u>C</u>hromatin <u>ac</u>cessibility and transcriptomics <u>u</u>nifying <u>s</u>oftware), a pipeline coded in Nextflow for cache-friendly, efficient, and scalable deployment, with all tools encapsulated in containers (Docker or Singularity) or in virtual environments (conda or Mamba [15]) for hassle-free installation and high reproducibility (Fig. 1a). Cactus is a versatile software that can be employed to analyze mRNA-Seq or ATAC-Seq data separately or jointly, allowing for the investigation of interrelated changes in chromatin accessibility and gene expression. The pipeline preprocesses data, performs differential analysis, and splits results into <u>D</u>ifferential <u>A</u>nalysis <u>S</u>ubsets (DASs). Enrichment analysis is then conducted on each DAS using several internal and external datasets, providing comprehensive molecular insights (Fig. 1b-d).

**Figure 1.**
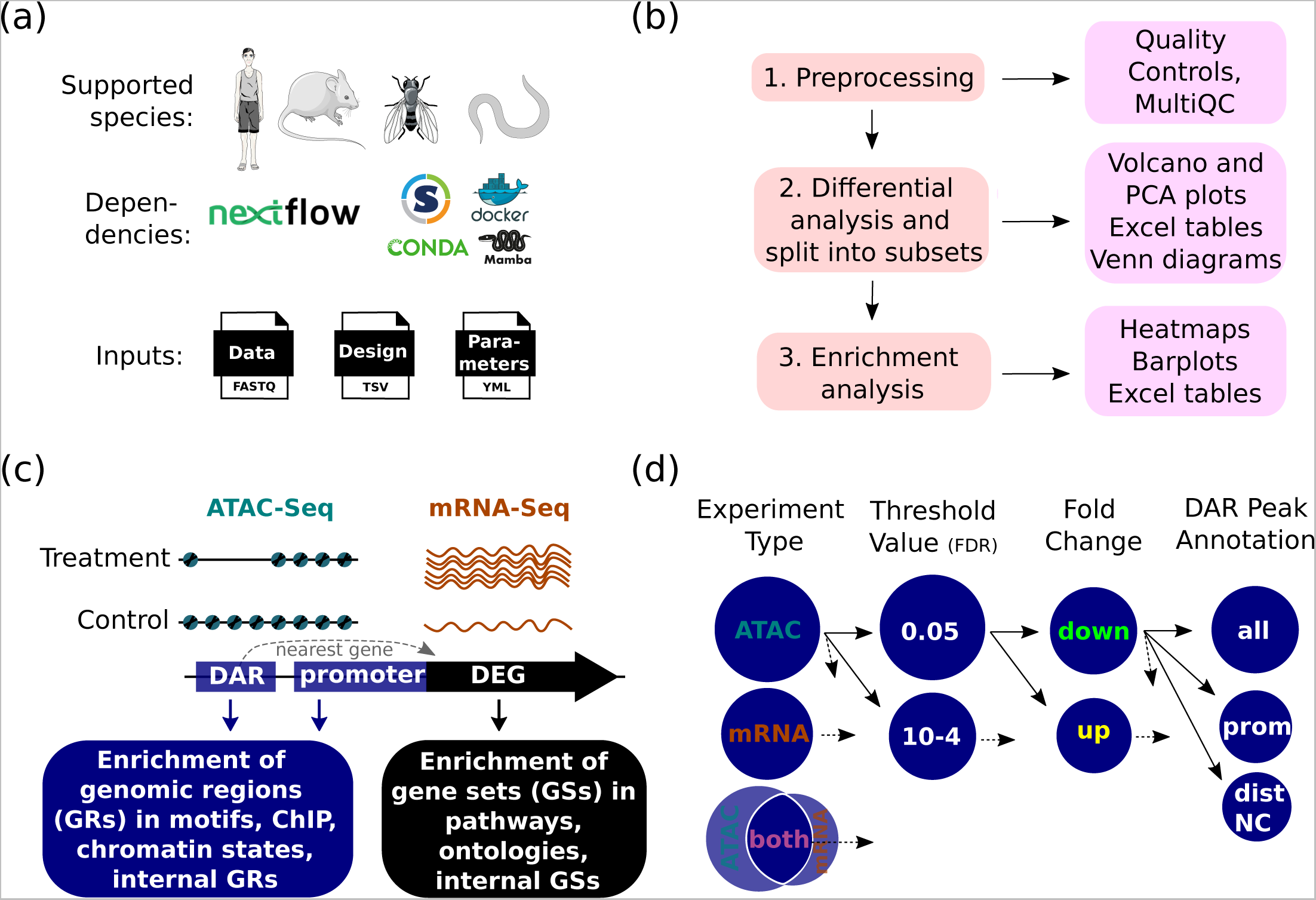
Overview of Cactus. (a) Key features. Species icons were adapted from [79,80]. (b) Simplified workflow. (c) Example of enrichment analysis performed for a gene showing an increase in both chromatin accessibility and gene expression upon treatment. Enrichment of internal genomic regions (GRs) and gene sets (GSs) indicates enrichment of GRs and GSs in other GRs and GSs generated by the pipeline. Black lines and blue circles represent DNA and nucleosomes, respectively. Orange lines represent mRNA molecules. (d) Sub-workflow showing the creation of DASs. Dotted arrows indicate optional additional filters. Abbreviations: DAR, <u>d</u>ifferentially <u>a</u>ccessible region; DEG, <u>d</u>ifferentially <u>e</u>xpressed <u>g</u>ene; ChIP, ChIP-Seq binding sites; motifs, DNA binding motifs; FDR, false <u>d</u>iscovery rate; prom, <u>prom</u>oter; distNC, <u>dist</u>al <u>n</u>on-<u>c</u>oding region.

We tested Cactus by running it on ATAC-Seq and mRNA-Seq data generated from *Caenorhabditis elegans* whole animals and human cells. To illustrate Cactus’ capabilities, we provide examples of outputs generated during the *C. elegans* case study analysis in Fig. 2 and 3, and the *C. elegans* test dataset analysis in Fig. S1, S2 and S3. The biological interpretation of the case study analysis is presented in the subsequent figures (Fig. 3 to 7). Importantly, we showed that Cactus could recapitulate the main findings of the original study and also provide some novel findings. In particular, Cactus successfully identified the key roles played by FOS and Polycomb-Repressive Complexes (PRCs) in regulating reprogramming, which resonated with recently published studies [16–24]. Furthermore, Cactus identified interesting new candidates regulating reprogramming and delineated their effects in different tissues.

**Figure 2.**
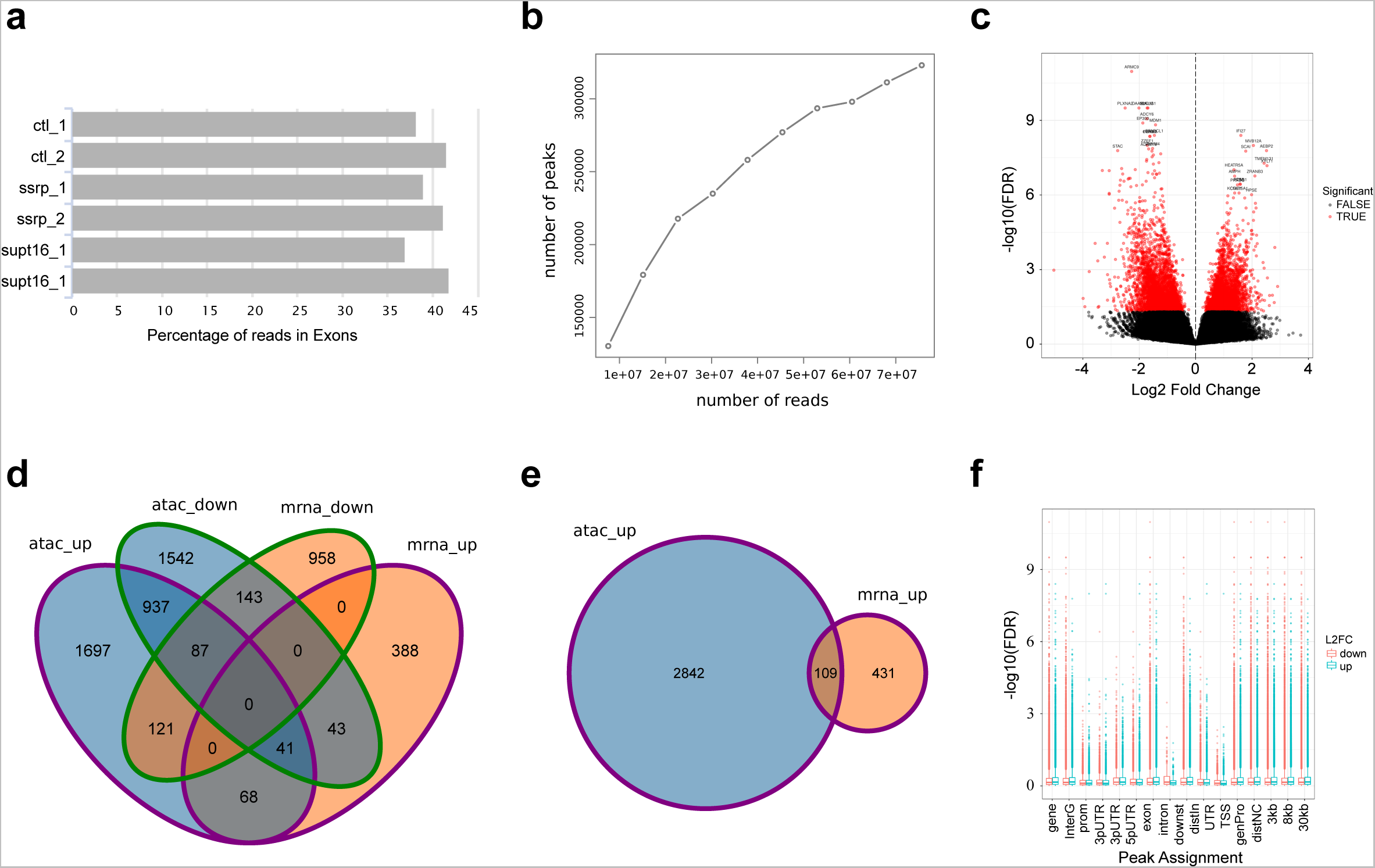
Example of outputs for the preprocessing and differential analysis steps. (a) Percentage of reads aligned to exons for 6 samples. (b) Saturation analysis for one replicate of the control condition. Panels (c-f) show figures for the comparison SSRP1 knockdown vs control. (c) Volcano plot. (d) Four-way Venn diagram showing the overlap between DEG- and DAR-associated genes. Please observe that multiple DARs of opening and closing chromatin can be associated to the same gene which explains the non-empty overlaps of the atac_up and atac_down sets. (e) Proportional two-way Venn diagram showing the overlap between mRNA-Seq and ATAC-Seq DASs. (f) Significance level of DARs assigned to each of the 19 available DAR Peak Annotation filters. Abbreviations: ctl, <u>c</u>ontrol.

**Figure 3.**
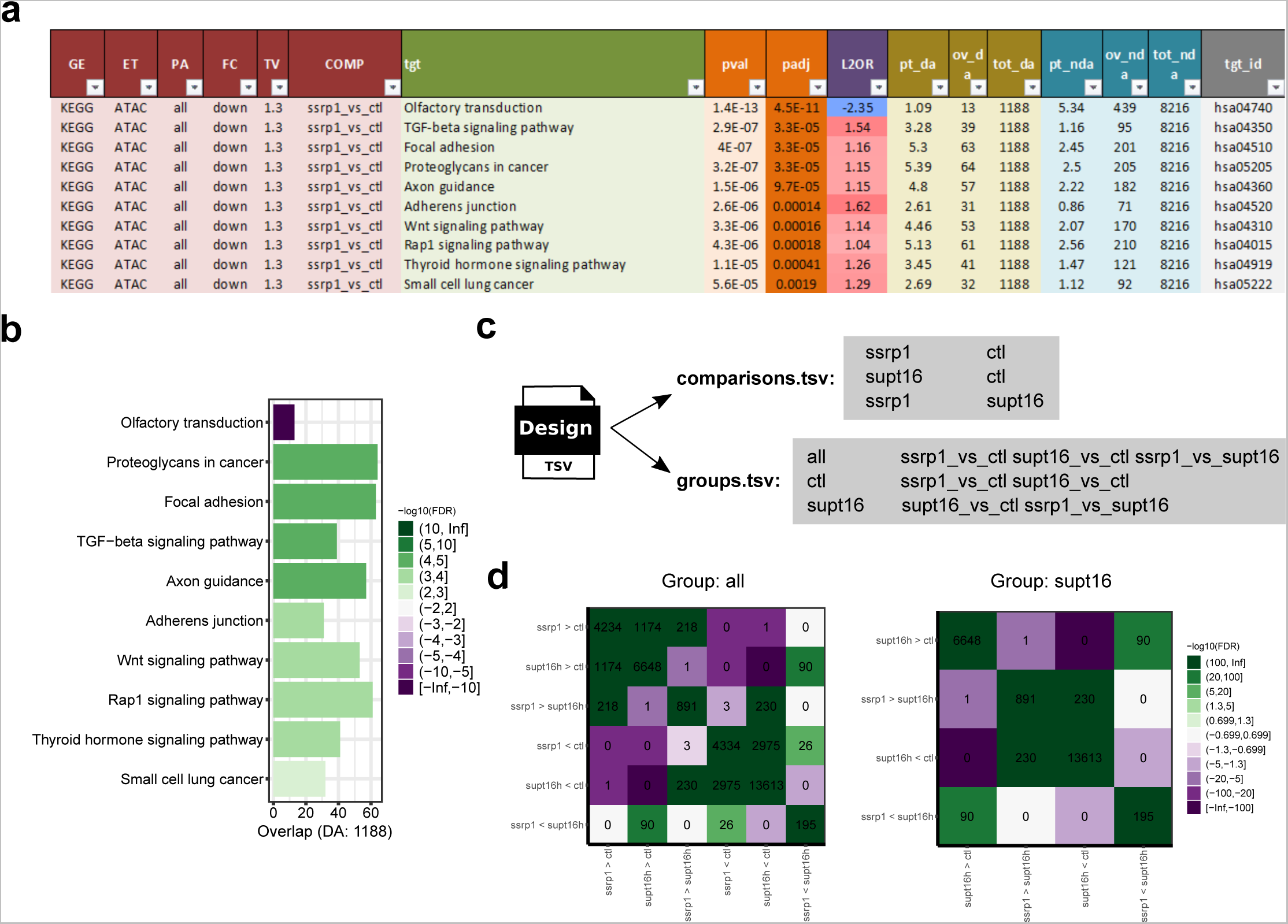
Example of outputs for the enrichment analysis steps. Panels (a-b) show KEGG enrichment analysis results for the comparison SSRP1 knockdown vs control. (a) Formatted Excel table. (b) Barplot. (c) Content of the TSV file used to define the comparisons to make between conditions and the groups of comparisons to plot together in heatmaps. (d) Heatmaps of the enrichment between comparisons for the “all” group and the “supt16h” group defined in (c). The “ctl” group is shown in Figure 5d.

## Material and Methods

### 1. Creation of references

All the references required by Cactus are parsed by an efficient Nextflow (v22.10.8) script that uses containers to improve ease of use, speed, and reproducibility. Currently, Cactus supports the four ENCODE/modENCODE species: *Homo sapiens*, *Mus musculus*, *Drosophila melanogaster,* and *C. elegans* [25–28]. Genomic sequences and annotations were downloaded from Ensembl [29], using the Ensembl release 107 for assemblies ce11, dm6 (BDGP6.32) and hg38, and the Ensembl release 102 for assembly mm10. An older Ensembl release was used for *M. musculus* as mm39 was not yet available for HOMER [30] files and Encode bed files. The ENCODE API [31] was used to download the ChIP-Seq binding sites of 2,714 Transcription Factors (TFs) [32,33] and chromatin states in the form of 899 ChromHMM profiles [34,35] and 6 HiHMM profiles [36]. Slim annotations (cell, organ, development, and system) were parsed and used to create groups of ChIP-Seq binding sites that share the same annotations, allowing users to analyze only ChIP-Seq binding sites relevant to their study. 2,779 TF DNA binding motifs were obtained from the Cis-BP database (build 2.00) [37]. The R packages AnnotationHub (v3.2.0) [38] and clusterProfiler (v4.8.1) [39,40] were used to obtain GO terms (by downloading NCBI orgdb v3.14) and KEGG pathways, respectively. HOMER files were downloaded for genomes (v6.4), organisms (v6.3), and promoters (v5.5).

### 2. Data analysis workflow

The three main analysis steps of Cactus are 1. Preprocessing, 2. Differential Analysis, and 3. Enrichment analysis. A large part of the pipeline consists in conducting preprocessing and quality controls on ATAC-Seq input samples. This can be seen in the detailed flowchart of the pipeline, available in Fig. S4, in which the connections between all Nextflow processes are displayed. Below, we provide details on the three main analysis steps.

#### 2.1 Preprocessing

Quality controls on raw reads are performed using FastQC (v0.11.7) [41] and are presented in multiQC reports (v1.18) [42] for both ATAC-Seq and mRNA-Seq data. For ATAC-Seq, reads are trimmed with skewer (v0.2.2) [43], compressed in parallel using pigz (v2.6), and aligned with Bowtie2 (v2.4.4) [44]. Randomly sampled raw and aligned reads are used to rapidly generate various quality control figures that are included in the multiQC report (Fig. 2a). Low-quality, duplicated, and mitochondrial reads are removed, and insert size distributions are plotted (Fig. S1a). Bigwig tracks, coverage (Fig. S1b) and PCA (Fig. S1c) plots, as well as correlations heatmaps between samples (Fig. S1d) are generated using deepTools (v3.4.3) [45]. Aligned reads are converted to bed format and shifted to account for the shift of the transposase [9]. Peaks are called using MACS2 (v2.2.7.1) [46]. Saturation curves are generated to indicate if re-sequencing is necessary (Fig. 2b, S1e). Peaks are split into sub-peaks and filtered for blacklisted regions, genomic DNA (i.e., input control) peaks (if available), and custom hotspot regions if specified. Then, ATAC-Seq peaks are annotated with ChIPseeker (v1.36.0) [47] and summary figures are made with clusterProfiler (v4.8.1) [39,40] (Fig. S1f-j). For mRNA-Seq, gene expression levels are quantified using kallisto (v0.46.2) [48].

#### 2.2 Differential analysis

For ATAC-Seq data, <u>d</u>ifferentially <u>a</u>ccessible regions (DARs) are estimated by DiffBind (v3.4.0) [49] and these regions are then annotated to the closest gene with ChIPseeker (Fig. 1c). For mRNA-Seq data, <u>d</u>ifferentially <u>e</u>xpressed <u>g</u>enes (DEGs) are determined with sleuth (v.0.30.0) [50]. Volcano (Fig. 2c, S2a,f) and PCA (Fig. S2b,c,g,h) plots, Venn diagrams (Fig. 2d,e, S2m-o), as well as standardized tables (Fig. S2p-v) are generated. Differential analysis results are then split into subsets (i.e., DASs) (Fig. 1d, S5) to perform enrichment analyses from various angles and extract the most insights out of the data. For instance, certain TFs might be binding only in intergenic regions. The enrichment of their motifs could therefore be masked when analyzing all peaks if most peaks are not in intergenic regions. Similarly, certain TFs may be strongly activating a small set of target genes, and therefore the enrichment of their motifs could be masked when considering all significantly activated genes.

Filters used to split data are: experiment type (ATAC, mRNA, or both; see Fig. S5), fold change type (up or down), significance level (adjusted p-value cutoff or top N entries), and DAR Peak Annotation (PA) to genomic region (currently 19 options, such as: all, promoter, gene, intergenic, etc.). Experiment type “both” refers to low chromatin <u>a</u>ccessibility and low <u>e</u>xpression (LA-LE) as well as <u>h</u>igh chromatin <u>a</u>ccessibility and <u>h</u>igh <u>e</u>xpression (HA-HE) genes (see examples in Fig. S6). The significance level of DARs by PA filter is plotted to help identify PA filters to use for downstream enrichment analysis (Fig. 2f, S2l).

#### 2.3 Enrichment analysis

For each DAS, Cactus performs enrichment analyses among experimental conditions as well as with external databases of gene ontologies, pathways, DNA binding motifs, ChIP-Seq binding sites, and chromatin states (Fig. 1c). For all enrichment analyses, the overlaps of the target (e.g., a pathway) with the DAS and the remaining entries (differential analysis results not in the DAS) are computed. Overlaps are computed using clusterProfiler for functional annotations and HOMER (v4.9.1) [30] for TF DNA binding motifs. We employed custom scripts for other enrichment categories, such as pathways, ontologies, chromatin states or DASs. Then, two-sided Fisher’s exact tests are conducted to identify enriched and depleted entries [51], and corrections for multiple testing are done using Benjamini and Hochberg’s False Discovery Rate (FDR) [52]. Results are presented in barplots (Fig. 3b, S3a-g), heatmaps (Fig. 3d, S3h-n) and tables (Fig. 3a, S3o-u). Heatmaps, which display DASs that pass the same filters (except the Fold Change filter), are particularly useful for quickly identifying patterns in complex experimental designs. They can be customized by specifying user-defined groups of comparisons to plot together (Fig. 3c) and by multiple options (e.g., adding dots proportional to log2 odds ratios or overlap size, plotting both enrichment and depletion or enrichment only, etc.).

### 3. Implementation

Cactus is a computational tool primarily written in the Nextflow, shell, and R programming languages. Most of the tools utilized by Cactus are packaged within Biocontainers [53,54], which ensures consistency across container images and virtual environments. One of the key advantages of Cactus is its ability to run offline. This feature ensures stable runtime performance and allows for the analysis of sensitive patient data without any security concerns. Installation is user-friendly, as the necessary containers or virtual environments are automatically downloaded the first time the pipeline is run. To help users become familiar with the pipeline, test datasets for each species have been created by downloading publicly available datasets with fetchngs (v1.6) [14] and subsampling the reads to the minimum required for running the pipeline. Test datasets can be run using one-liners as shown in the online documentation (in the Quick Start and the Tutorial sections). These test datasets and references are available on Figshare [55]. Before using Cactus, users should download the necessary references and, optionally, test datasets. Cactus can be executed by providing it with a YML file that specifies the run parameters, including the location of TSV files outlining the experimental design and the path to the raw FASTQ.gz data files (Fig. 1a, 3c).

## Results and discussion

### Comparisons with existing tools

We set out to compare Cactus with recently published ATAC-Seq analysis tools (Table 1). We found that while several tools provide motif enrichment analysis, this analysis step is almost always done on raw peaks from a single condition. However, biological experiments typically involve comparison of a given condition to another condition, such as a control. Therefore, enrichment analysis is more informative when performed on entries selected from differential analysis. To our knowledge, CoBRA [56] is the only published pipeline, besides Cactus, to provide this important functionality. Furthermore, Cactus provides several unique functionalities not provided by any other tools such as splitting DA results into subsets, enrichment in chromatin states and ChIP-Seq binding sites, creation of customized heatmaps, and the reporting of results in merged pdfs or tables.

**Table 1.** Comparison of the main features and implementation of recently published ATAC-Seq pipeline tools. Abbreviations: DA, Differential Analysis.

| Category | Feature | esAT<br>AC | ATAC-pipe | Alfre<br>d | snake<br>Pipes | nfc<br>or<br>e/ata<br>c | ATAC<br>graph | AIAP | CoBR<br>A | PEPA<br>TAC | Cactu<br>s |
| --- | --- | --- | --- | --- | --- | --- | --- | --- | --- | --- | --- |
| Pipeline architecture | Workflow manager | — | — | — | Snake<br>make | Nextfl<br>ow | — | — | Snake<br>make | Pypip<br>er | Nextfl<br>ow |
|  | Containers | — | — | ✓ | — | ✓ | ✓ | ✓ | ✓ | ✓ | ✓ |
|  | Virtual environments | — | — | ✓ | ✓ | ✓ | — | — | ✓ | ✓ | ✓ |
|  | Built-in pipeline(s) | ✓ | ✓ | — | — | ✓ | ✓ | ✓ | ✓ | ✓ | ✓ |
|  | Quality controls on reads | ✓ | ✓ | ✓ | ✓ | ✓ | ✓ | ✓ | — | ✓ | ✓ |
| Preprocessing ATAC-Seq data | Peak annotation | ✓ | — | ✓ | ✓ | ✓ | ✓ | ✓ | ✓ | ✓ | ✓ |
|  | Saturation analysis | — | — | — | — | — | — | ✓ | — | — | ✓ |
|  | Enrichment of motifs in raw ATAC-Seq peaks | ✓ | ✓ | ✓ | — | ✓ | — | — | — | ✓ | — |
|  | Differential accessibility analysis | — | ✓ | — | ✓ | ✓ | ✓ | ✓ | ✓ | — | ✓ |
| Differential analysis | Venn diagrams | — | — | — | — | — | ✓ | — | — | — | ✓ |
|  | Volcano plots | — | ✓ | — | — | — | — | — | ✓ | — | ✓ |
|  | Integration with mRNA-seq data | — | — | — | — | — | ✓ | — | — | — | ✓ |
|  | Splitting DA results into subsets | — | — | — | — | — | — | — | — | — | ✓ |
| Enrichment analysis for DARs (or DASS) | Enrichment of motifs | — | — | — | — | — | — | — | ✓ | — | ✓ |
|  | Enrichment of ChIP-Seq binding sites | — | — | — | — | — | — | — | — | — | ✓ |
|  | Enrichment of chromatin states | — | — | — | — | — | — | — | — | — | ✓ |
|  | Enrichment of ontologies and pathways | — | — | — | — | — | — | — | ✓ | — | ✓ |
|  | Enrichment of comparisons | — | — | — | — | — | — | — | —<br>(clustering) | — | ✓ |
|  | Creation of customized heatmaps | — | — | — | — | — | — | — | — | — | ✓ |
| Reporting | Reporting results in merged pdfs or tables | — | — | — | — | — | — | — | — | — | ✓ |
|  | Optional genomic DNA control | ✓ | — | — | — | — | ✓ | — | — | — | ✓ |
|  | Test dataset(s) | ✓ | ✓ | — | ✓ | — | — | ✓ | ✓ | ✓ | ✓ |
|  | Automatic download of references | ✓ | ✓ | — | ✓ | — | — | — | ✓ | — | ✓ |
| Year |  | 2018 | 2019 | 2019 | 2020 | 2020 | 2021 | 2021 | 2021 | 2021 | 2024 |
| Reference |  | [57] | [58] | [59] | [60] | [14] | [61] | [62] | [56] | [63] | — |

### Outputs files

The outputs of Cactus are organized in 5 directories: Figures_Individual, Figures_Merged, Tables_Individual, Tables_Merged, and Processed_data; each of which contains the following folders: 1_Preprocessing, 2_Differential_Analysis, and 3_Enrichment analysis. Cactus outputs contain figures for the pre-processing of reads (e.g., insert size, correlation, PCA) and peaks (e.g., average profile, coverage, saturation curve), differential analysis (e.g., PCA and volcano plots, Venn diagrams), and enrichment analysis (e.g., barplots and heatmaps for the various enrichment categories). Produced tables include detailed and filtered tables for ATAC-Seq, mRNA-Seq and DASs. Processed data include reads, peaks, and quality controls at various steps of the analysis as well as R objects (e.g., gene sets, DiffBind, ChIPseeker, sleuth, GRange, and ggplot2 objects), and genomic regions for results of differential and enrichment analysis.

## Case study

### Introduction to the original study: Identification of FACT as a complex regulating reprogramming in C. elegans and human samples

To demonstrate the utility of Cactus, we reanalyzed data from a study by the Tursun laboratory, in which a histone chaperone complex, called FAcilitates Chromatin Transcription (FACT), was identified as a cellular reprogramming barrier in *C. elegans* [64]. The authors overexpressed CHE-1, a C2H2 zinc finger Transcription Factor (TF) that is essential for specifying ASE gustatory neurons [65]. Overexpression of CHE-1 alone is insufficient to induce the reprogramming of non-neuronal cells into ASE neurons. Therefore, the authors screened a whole-genome RNA interference library for knockdowns that would overcome this reprogramming barrier and induce expression of an ASE neuronal reporter gene. The most important hits from this screen were *C. elegans* orthologs of FACT subunits (HMG-3, HMG-4, SPT-16) and genes known to functionally interact with FACT in other species. Interestingly, *C. elegans* orthologs of FACT subunits showed tissue-specific effects, with HMG-3 and SPT-16 acting as barriers to reprogramming in germ cells, while HMG-4 and SPT-16 acted as barriers in the intestine. This discovery was further reinforced by the subsequent finding that human FACT subunits, SSRP1 and SUPT16H, acted as barriers for reprogramming human fibroblasts into induced pluripotent stem cells.

### ATAC-Seq and mRNA-Seq results described in the original study

To investigate the molecular mechanisms behind these reprogramming barriers, the authors conducted ATAC-Seq and mRNA-Seq experiments under normal or reduced levels of FACT subunits in both *C. elegans* and human fibroblasts. The results showed a strong correlation between chromatin accessibility and transcriptional changes caused by the depletion of FACT subunits in both organisms. Even though FACT was previously known to activate transcription [66], the authors found an equal number of chromatin regions opening and closing and observed equal numbers of gene induction and repression events. Moreover, they observed expression changes in genes involved in cellular reprogramming in human fibroblasts, with nine reprogramming-preventive genes being downregulated and eight reprogramming-promotive genes, including CEBP (CCAAT/Enhancer-Binding Protein), being upregulated upon FACT depletion. The authors conducted a *de novo* motif analysis on the FACT-dependent ATAC-Seq peaks to identify TFs that may bind these regions and hence act downstream. Upon FACT depletion in *C. elegans,* binding motifs of ELT-2/7, CEH-20/40, and UNC-62 were enriched in closing chromatin regions, while binding motifs of JUN-1 were enriched in opening chromatin regions. In human fibroblasts, the binding motif of RUNX1/3 was enriched in closing regions, while the binding motifs of CEBPA, CEBPE, JUN, and FOSL2 were enriched in opening regions. Interestingly, both JUN and CEBP TFs were previously shown to promote reprogramming in *C. elegans* and human cells, respectively. The authors concluded that FACT acts as both an activator and a repressor of gene expression, and that JUN and CEBP may play important roles in the FACT-dependent repression of reprogramming-promoting genes.

### Cactus could recapitulate the main findings of the original study

We ran Cactus on *Kolundzic et al.*’s data using standard parameters. Thus, all results shown below represent differential analysis subsets (DASs) with adjusted p-value cutoffs of 0.05. Our reanalysis confirmed the strong correlation between changes in chromatin accessibility and changes in transcription following knockdown of FACT subunits (Fig. 4, False Discovery Rate < 10^-100^ for all comparisons). We also observed a similar number of up- and down-regulated genes in both organisms (Fig. 4a,b), thereby confirming that FACT has both gene-activatory and -repressive effects. We then specifically examined the effect of FACT depletion on the transcription of key reprogramming genes in human fibroblasts, as defined in Figure 6L of *Kolundzic et al.* Looking at the direction of change in expression for 16 genes (7 reprogramming-promotive genes and 9 reprogramming-preventive genes) upon knockdown of FACT subunits we obtained results consistent with those of the original study for 29 out of 32 entries (Table S1). This confirmed the regulation of key reprogramming genes by the FACT complex. Lastly, our analysis confirmed most of the TF binding motif enrichments in regions opened or closed by FACT, namely enrichments for ELT-2/7, JUN-1, and RUNX1/3. Only the enrichments of UNC-62 and CEH-20/40 binding motifs in *C. elegans* could not be recapitulated (Fig. 5a,c). We conclude that our Cactus analysis was able to recapitulate all major ATAC-Seq- and mRNA-Seq-derived results and conclusions of the original study.

**Figure 4.**
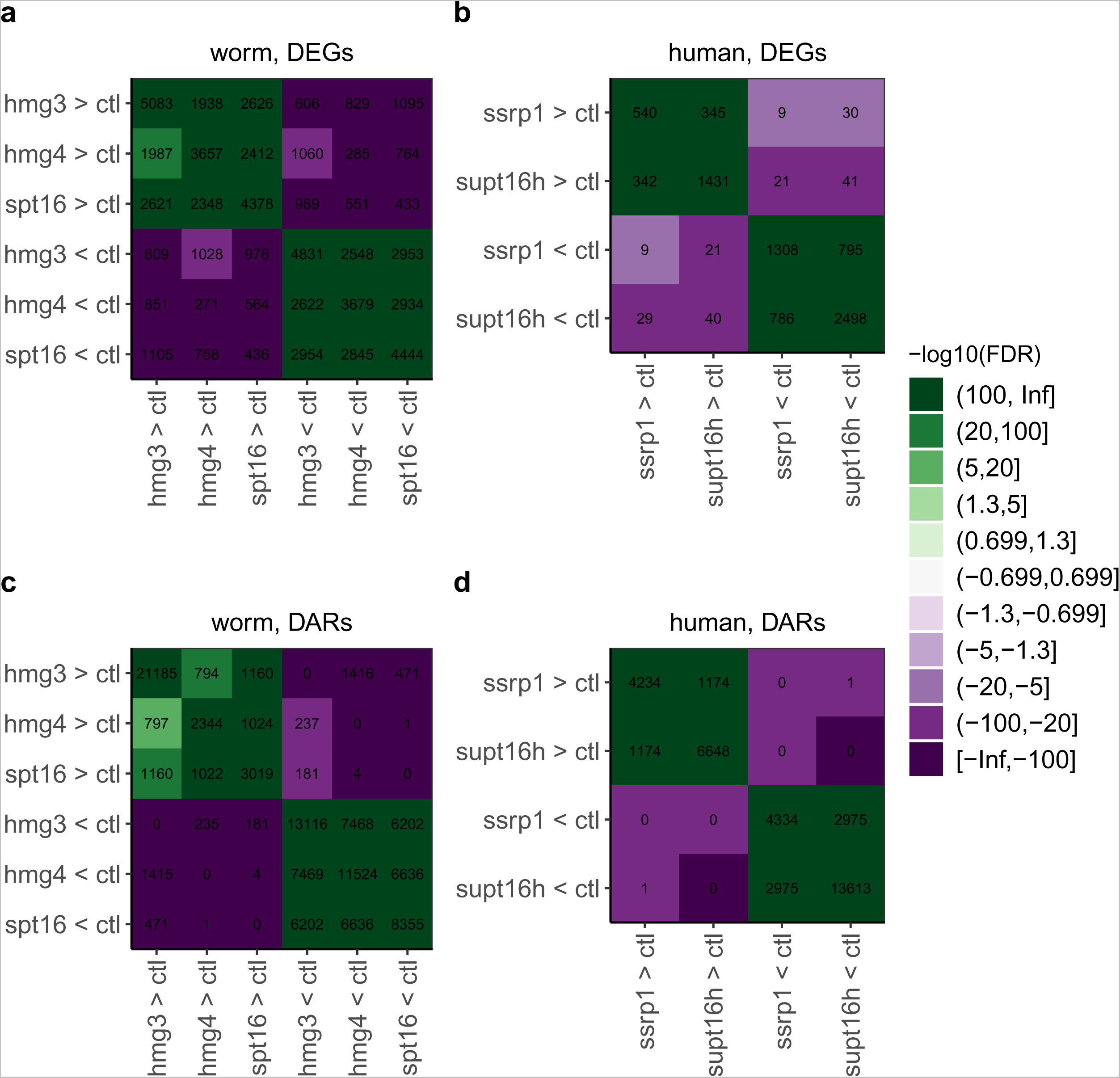
Enrichment of DASs in other DASs. Heatmaps showing the overlap between DEGs from mRNA-Seq (a,b) and DARs from ATAC-Seq (c,d) for *C. elegans* (a,c) and human (b,d) samples. Numbers and colors indicate the size and significance of the overlap, respectively. Colors show the signed minus log_10_ adjusted p-values (from two-sided Fisher tests), with positive values (green color) indicating enrichment and negative values (purple color) indicating depletion. The (0.699, 1.3] and (100, Inf] bins correspond to the adjusted p-value intervals of (0.2, 0.05] and (10^-100^, 0], respectively. Abbreviation: ctl, <u>c</u>ontrol.

**Figure 5.**
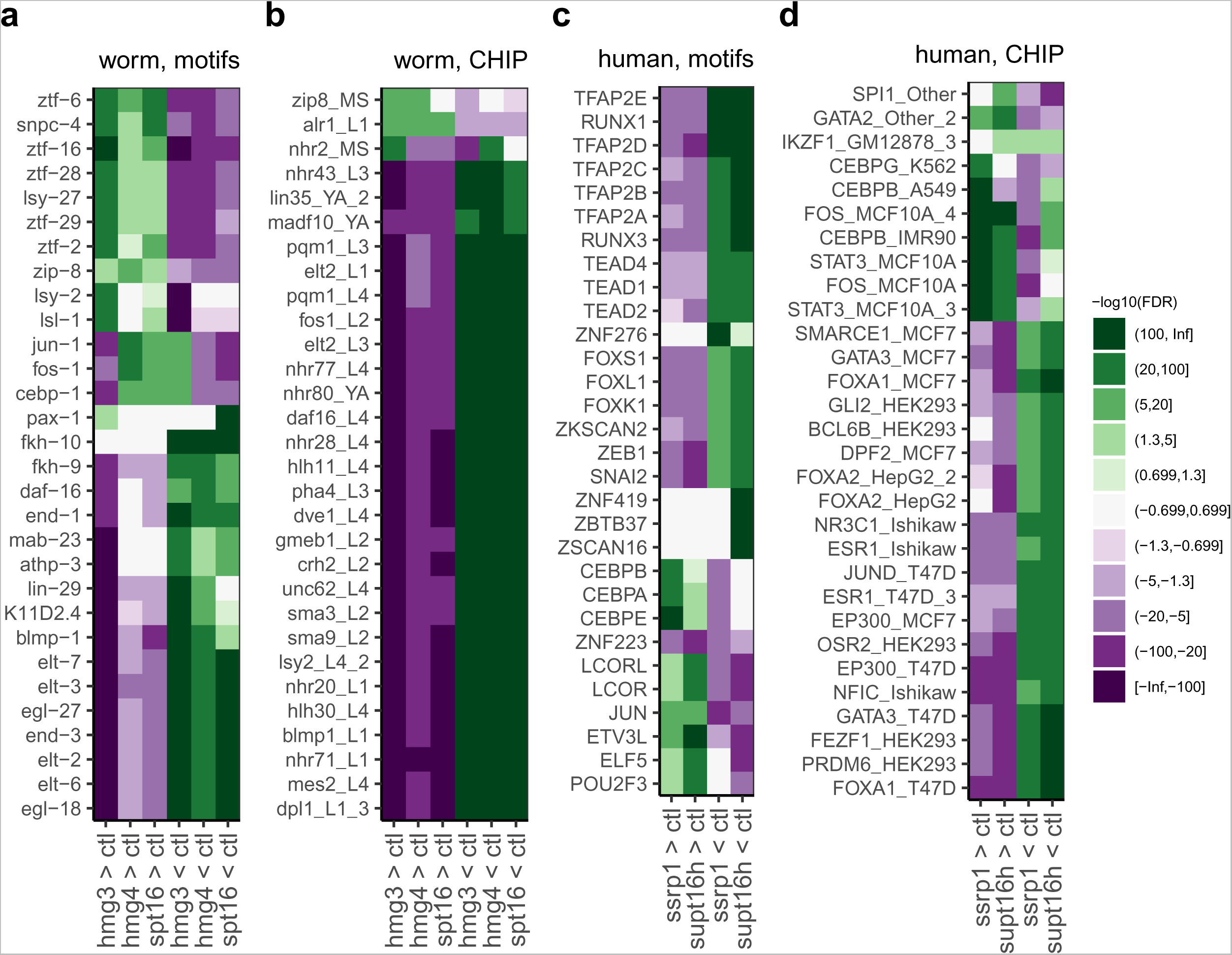
Enrichment of TFs CHIP-Seq binding sites and DNA binding motifs in DARs. Colors indicate the signed minus log_10_ adjusted p-values (from two-sided Fisher tests), with positive values (green color) indicating enrichment and negative values (purple color) indicating depletion. The (0.699, 1.3] and (100, Inf] bins correspond to the adjusted p-value intervals of (0.2, 0.05] and (10^-100^, 0], respectively. For (b) and (d), the y-axis labels are in the following format: protein_*C. elegans* developmental stage (b) or protein_cell line (d). Details about the cell lines in which ChIP-Seq experiments were conducted are available in the reference section of the Cactus documentation. Abbreviation: ctl, <u>c</u>ontrol.

**Figure 6.**
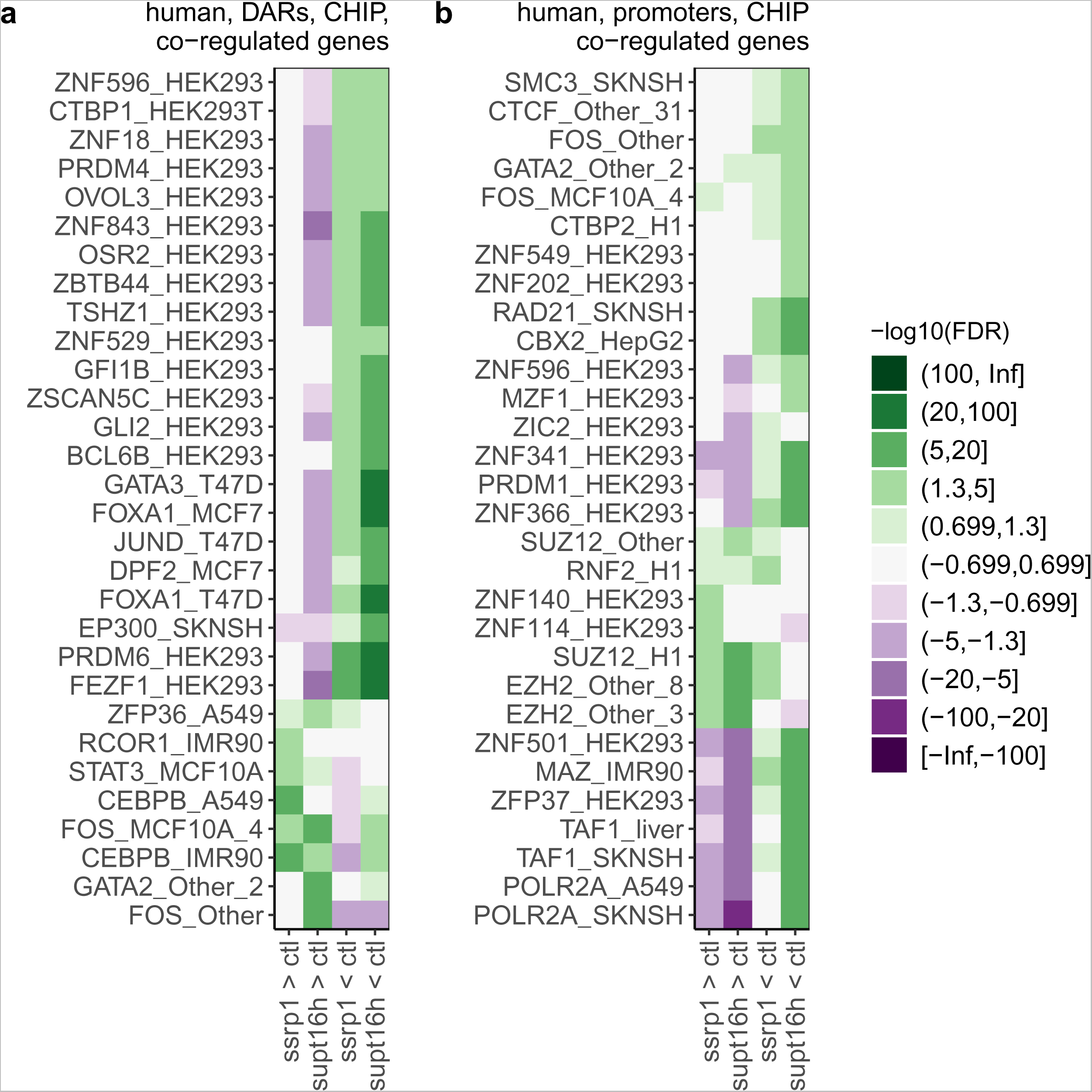
Enrichment of TFs CHIP-Seq binding sites in DASs from HA-HE and LA-LE genes. Enrichment of DARs (a) and promoters (b) of HA-HE (High chromatin Accessibility and High Expression) and LA-LE (Low chromatin Accessibility and Low Expression) genes upon FACT knockdown in TFs CHIP-Seq binding sites. Colors indicate the signed minus log_10_ adjusted p-values (from two-sided Fisher tests), with positive values (green color) indicating enrichment and negative values (purple color) indicating depletion. The (0.699, 1.3] and (100, Inf] bins correspond to the adjusted p-value intervals of (0.2, 0.05] and (10^-100^, 0], respectively. The y-axis labels are in the following format: Protein_Cell line. Details about the cell lines in which ChIP-Seq experiments were conducted are available in the reference section of the Cactus documentation. Abbreviation: ctl, <u>c</u>ontrol.

### Cactus could confidently predict a role of FOS in regulating reprogramming

Notably, the advanced analysis pipeline of Cactus allowed us to make several additional observations not described in the original study. For example, we obtained additional evidence that CEBP and JUN/FOS bind to regions opening upon FACT depletion. Kolundzic *et al.* had shown that the DNA binding motif of CEBP-1 was highly enriched in FACT-dependent regions of human cells. While Cactus reproduces this observation, it also finds the same for *C. elegans* (Fig. 5a). Additionally, Cactus found ChIP-Seq binding sites of both CEBP and FOS (a binding partner of JUN, together forming the AP-1 complex) highly enriched in FACT-dependent regions (Fig 5d). In Cactus, genes with increased chromatin accessibility and increased gene expression in one condition versus another (i.e., FACT versus wild-type control) are called High chromatin Accessibility and High Expression (HA-HE) genes [67] (Fig. S5). Strikingly, we found that FOS and CEBP were the most enriched ChIP-Seq binding sites in ATAC-Seq peaks assigned to HA-HE genes (Fig. 6a). Taken together, our observations strongly suggest that FACT represses regions that are bound by JUN/FOS and CEBP. Kolundzic *et al.* reported the enrichment of the FOSL2 motif in chromatin that opens upon SUPT16H depletion. However, this result was only reported in one supplementary figure, for one organism and one FACT subunit. In contrast, Cactus could robustly detect such pattern in both species and for all FACT subunits. This result was even more striking when jointly analyzing ATAC-Seq and mRNA-Seq data. Interestingly, recent reports have confirmed that FOS plays a key role in the regulation of reprogramming [16–19], supporting the results from our analysis.

### Cactus could identify interesting TF candidates that may cooperate with FACT to regulate reprogramming in multiple tissues

In their study, *Kolundzic et al.* provided several ATAC-Seq and mRNA-Seq datasets related to FACT and their analysis, including RAW data generated upon knockdown of the germline-specific FACT subunit HMG-3 in *C. elegans*. However, their combined analysis was missing. We now analyzed this data and found that HMG-3 depletion results in similar chromatin accessibility and gene expression changes as the depletion of other FACT subunits, e.g., HMG-4 and SPT-16, with the exception of an additional ∼20,000 regions where chromatin opened only upon HMG-3 depletion (Fig. 4c). Unexpectedly, we found that the JUN/FOS and CEBP-1 DNA binding motifs were highly depleted in these regions (Fig. 5a), implying that JUN/FOS and CEBP-1 show tissue-specific differences when partnering with FACT. This raises the question of whether FACT partners with other TFs in both the gut and the germline.

In *C. elegans*, the DNA-binding motifs (Fig. 5a) or ChIP-Seq binding sites (Fig. 5b) of several TFs involved in neuronal development (ALR-1, ZTF-16, ZTF-6, LSY-27) were enriched in chromatin regions that opened upon FACT depletion. Notably, of these, LSY-27 is involved in regulating the asymmetry of ASE neurons [68], which are the neurons specified by CHE-1 overexpression. Similarly, the DNA binding motif of the forkhead transcription factor FKH-10 was highly enriched in chromatin regions closing upon FACT depletion (Fig. 5a), which resonates with recent reports showing that Forkhead TFs are versatile regulators of cellular reprogramming to pluripotency [69]. In human cells, Cactus identified not only FOS and CEBP but also STAT3 (Signal Transducer and Activator of Transcription 3) as TFs whose ChIP-Seq binding sites were highly enriched in chromatin regions opening upon FACT depletion (Fig. 5d). This is remarkable because STAT3 is a well-known promoter of reprogramming [70–72].

### Cactus could successfully predict a role of Polycomb-Repressive Complexes in the regulation of reprogramming

Our analysis of chromatin states showed that opened chromatin was enriched for Polycomb-repressed regions in the L3 larvae of *C. elegans* (Fig. 7a), in a human foreskin fibroblast primary cell line (Fig. 7b), and in a human lymphoblastoid cell line (Fig. 7c). In addition, we found EZH2, the core enzymatic subunit of the Polycomb Repressor Complex 2 (PRC2), SUZ12, another PRC2 subunit, and RNF2, a PRC1 subunit, to be the only TF ChIP-Seq binding sites enriched in the promoters of HA-HE genes upon depletion of either of the two FACT subunits in human cells (Fig. 6b), with the PRC2 subunits being the most strongly enriched. Altogether, these results indicate that the repressive function of FACT is achieved via Polycomb-mediated gene repression. Interestingly, a previous *C. elegans* study identified PRC2 as another barrier that hinders the conversion of germ cells to somatic neurons upon CHE-1 overexpression [73]. No connection was described between FACT and Polycomb in the Kolundzic *et al.* study. This result resonates with reports from multiple studies indicating a key role of PRC1 and PRC2 in the reprogramming process [20–24]. In conclusion, genes with high expression and high chromatin accessibility upon FACT depletion include key reprogramming-promoting genes and are located in Polycomb-repressed regions.

Our analysis showed that the DNA binding motif of SNPC-4 was highly enriched in chromatin opening upon FACT depletion in *C. elegans* (Fig. 5a). Interestingly, SNPC-4 is a component of the Upstream Sequence Transcription Complex (USTC) that binds to the H3K27-methylated promoter of piRNA genes, inducing their expression in a PRC2-dependent manner [74]. Furthermore, piRNAs have been recently shown to be differentially expressed specifically in reprogrammed stem cells and not in embryonic stem cells [75,76]. SNPC-4 has not been described previously as playing a role in reprogramming, and therefore, it could be an interesting candidate to evaluate for a potential role in the FACT- and PRC-2-mediated regulation of reprogramming.

### Conclusions of the case study

In summary, we successfully reproduced the main findings of the Kolundzic *et al.* study using Cactus. Our analysis confirmed the role of JUN and CEBP in reprogramming and identified new TF candidates such as FKH-10, LSY-27, and SNPC-4. Importantly, our joint analysis of ATAC-Seq and mRNA-Seq data allowed us to characterize genes with consistent changes in chromatin accessibility and expression levels upon FACT knockdown. This allowed us to discover that FACT prevents the opening of chromatin at PRC2-repressed promoters nearby regions bound by the reprogramming-promoting factors CEBP and FOS. These results are consistent with the key role of PRC2 and FOS for regulating reprogramming, as reported in recently published studies [16–24]. Finally, panels from Figures 2 to 7 were generated with just three calls to Cactus, making it a user-friendly and valuable tool for researchers to identify transcription factors that may be relevant to their conditions of interest.

**Figure 7.**
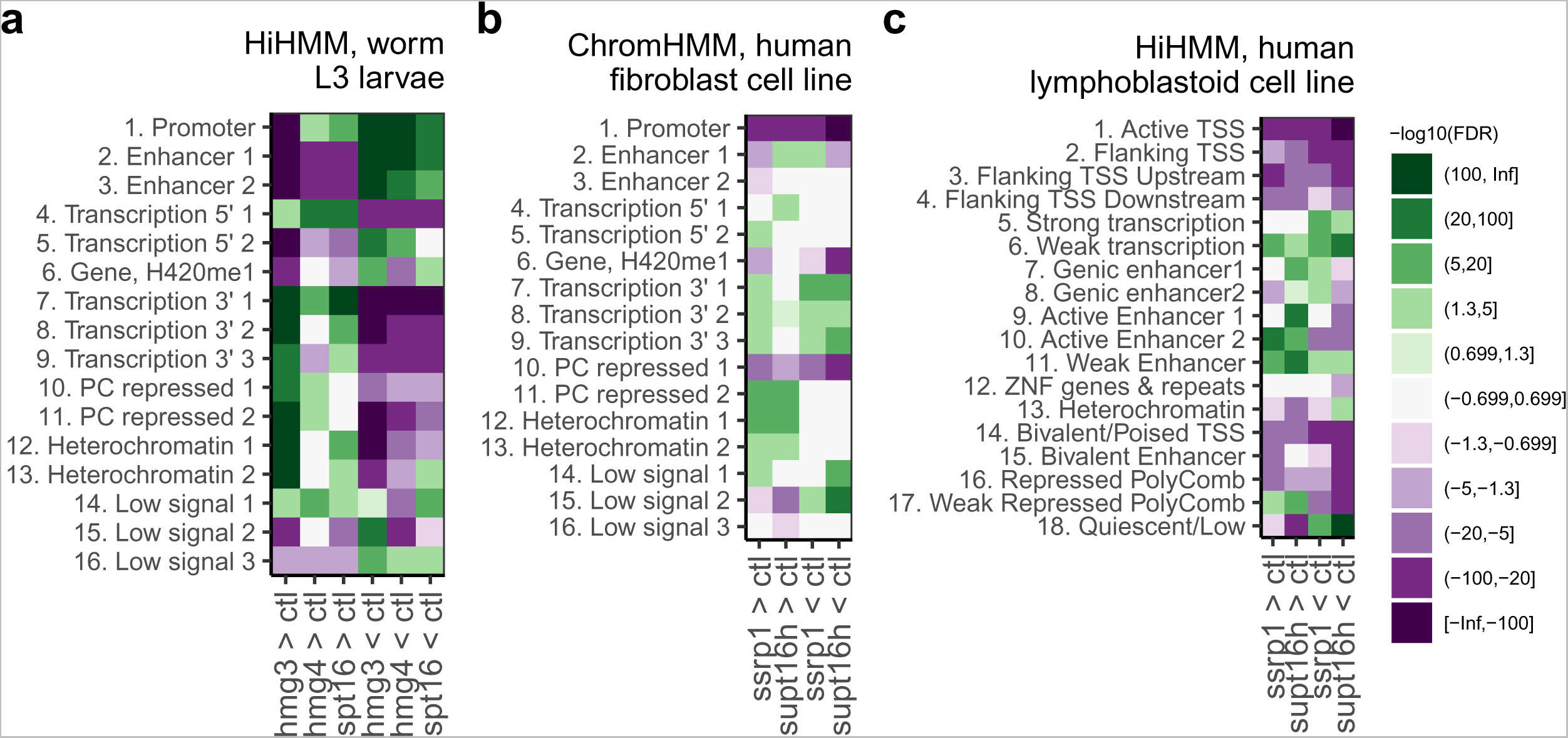
Enrichment of chromatin states in DARs. The chromatin state files used are a 17-state HiHMM model [36] for L3 *C. elegans* worms (a), a 17-state HiHMM model for B lymphocyte human cell line (b, cell line GM12878), and an 18-state ChromHMM model [34] for human primary foreskin fibroblast (c, ENCODE accession ENCSR251QDX). Colors indicate the signed minus log_10_ adjusted p-values (from two-sided Fisher tests), with positive values (green color) indicating enrichment and negative values (purple color) indicating depletion. The (0.699, 1.3] and (100, Inf] bins correspond to the adjusted p-value intervals of (0.2, 0.05] and (10^-100^, 0], respectively. Abbreviation: ctl, <u>c</u>ontrol; PC, <u>P</u>oly<u>c</u>omb.

## Limitations

The end-to-end capabilities of Cactus allow users with limited bioinformatics expertise to analyze their data in depth. However, the pipeline architecture currently has a limited flexibility in the choice of tools to use and the analysis steps that are performed. For instance, more advanced users may find it useful to perform batch correction or to use other differential analysis tools than the one used in Cactus. Nevertheless, it can be argued that advanced users may still run Cactus to effortlessly obtain a comprehensive overview of the patterns present in their data, and then develop analysis scripts customized to their dataset specificities that can recapitulate the interesting patterns they observed. Indeed, Cactus can streamline fine-grained enrichment analysis by splitting differential analysis results into various subsets, which helps in analyzing the data from various angles.

Another limitation of Cactus is that it supports only four commonly used species, which limits its use in other species. This design was adopted to leverage the comprehensive datasets generated by the ENCODE and modENCODE consortia, such as blacklisted regions, chromatin states, and ChIP-Seq binding sites. To remedy this, future versions of Cactus could include additional supported species, for which the use of blacklisted regions, as well as the analysis of chromatin states and ChIP-Seq binding sites, will be disabled.

## Conclusions

Cactus is a user-friendly and highly reproducible pipeline that allows researchers to perform in-depth analyses of their mRNA-Seq and/or ATAC-Seq datasets by comparing them internally and with various external datasets. The pipeline is designed for ease of use, with features such as the creation of formatted Excel tables, merged PDFs, and merged tables, while also allowing for a high degree of customization. We showed that Cactus stands out among similar tools by having several unique and important features, notably with its capacity to conduct various enrichment analyses on DASs and to report results in customizable heatmaps. When we applied Cactus to published datasets containing samples from *C. elegans* worms and human cells [64], our pipeline confirmed the main findings of the study, provided additional observations, and generated new molecular hypotheses, arguing that Cactus performs well in real-life scenarios. We anticipate that Cactus will be particularly useful for laboratories with limited time or bioinformatics expertise, and that it can help to reduce the reproducibility crisis seen in many omics studies [77] and science in general [78].

## Data availability

The data used in this article are available in the Gene Expression Omnibus (GEO) database at https://www.ncbi.nlm.nih.gov/geo under the GEO accession GSE98758. The Cactus code and documentation are available on Github at https://github.com/jsalignon/cactus as well as in the Zenodo open repository as a release archive at https://doi.org/10.5281/zenodo.11115632. Test datasets and references are available as a release archive in the Figshare SciLifeLab Data Repository at https://doi.org/10.17044/scilifelab.20171347.v4.

## Acknowledgement

We would like to thank Aaron C. Daugherty and the Brunet lab for sharing the code of their 2017 *C. elegans* ATAC-Seq manuscript, which was a valuable asset during the initial development of our ATAC-Seq analysis scripts. Additionally, we would like to thank all the scientists who have created the databases and tools used by Cactus, i.e., contributors to ENSEMBL, members of the ENCODE and modENCODE consortia, and creators of the Cis-BP database. Finally, we would like to thank Paolo Di Tommaso for providing technical support when developing Nextflow scripts.

## Author Contributions

J.S. and C.G.R. conceived the project. J.S. developed Cactus and all related scripts, created the test datasets, the references, the documentation, and conducted the case study analysis. L.M.A. provided initial test data and feedback on the design of Cactus. M.G. implemented support for Docker, conda, and Mamba, and provided guidance on the architecture of the pipeline. J.S. and C.G.R. wrote the manuscript.

