## Supplementary materials for "Cactus: a user-friendly and reproducible ATAC-Seq and mRNA-Seq analysis pipeline for data preprocessing, differential analysis, and enrichment analysis"

### Supplementary figure legends

Figure S1. Examples of figure outputs for the preprocessing part. (a) Insert size. (b) Read coverage. (c) PCA on reads. (d) Spearman correlation on reads. (e) Saturation curve. (f) Distribution of peaks in different types of genomic regions. (g) Average peak profiles of all samples. (h) Distribution of peaks’ distances to TSSs. (i) Average peaks profile of a sample. (j) Coverage of peaks across the genome. All figure panels originate from the *C. elegans* test dataset.

Figure S2. Examples of figure and table outputs for the differential analysis part. (a-e) Figures for differential gene expression analysis. (a) Volcano plot for mRNA-Seq. (b,c) PCA plots for mRNA-Seq showing PC3 and PC4 (b) and PC1 and PC2 (c). (d) Estimated counts density. (e) MA plot. (f-k) Figures for differential chromatin accessibility analysis. (f) Volcano plot for ATAC-Seq. (g,h) PCA plots for ATAC-Seq showing PC3 and PC4 (g) and PC1 and PC2 (h). (i) 4-way Venn diagram showing the first 2 replicates per condition. (j) Samples heatmap. (k) MA plot. (l) Boxplots of the significance of various Peak Annotation (PA) filters. (m-o) Venn diagrams. Blue background: ATAC-Seq. Orange background: mRNA-Seq. Purple lines: up-regulated. Green lines: down-regulated. (m) Proportional two-way Venn diagram for up- (m) and down- (n) regulated genes. (o) Four-ways Venn diagram. (p-r) Detailed ATAC-Seq Excel results table. (p) Main results. (q) Counts and annotated gene. (e) Peak annotation filters. (s-t) Detailed mRNA-Seq Excel results table. (s) Main results. (t) Sleuth’s estimates. (u) Synthetic ATAC-Seq and mRNA-Seq results table. (v) Results table containing all significant results from ATAC-Seq, mRNA-Seq, and their combination. All figure panels originate from the *C. elegans* test dataset.

Figure S3. Examples of figure and table outputs for the enrichment analysis part. (a-g) Barplots for the enrichment of ChIP-Seq peaks (a), chromatin states (b), functional annotations from GO-BP (Biological Processes) (c) and KEGG (d), DASs of genes (e), DASs of peaks (f), and TF DNA-binding motifs (g) in DASs. (h-m) Heatmaps for the enrichment of ChIP-Seq peaks (h), chromatin states (i), functional annotations from GO-BP (j) and KEGG (k), DASs of genes (l), TF DNA-binding motifs (m), and DASs of peaks (n) in DASs. (o-u) Excel results tables for ChIP-Seq peaks (o), chromatin states (p), GO-BP (q), KEGG (r), DASs of genes (s), TF DNA-binding motifs (t), and DASs of peaks (u). All figure panels originate from the *C. elegans* test dataset. Please observe that in (e) and (l) enrichment is calculated based on the overlap of gene sets (i.e., DEGs or genes associated to DARs), while in (f) and (n) it is calculated based on the overlap of genomic regions (i.e., DARs), as shown in Fig. 1c and S5.

Figure S4. Detailed workflow. The ellipses show the different analysis steps, with the utilized tool(s) written in italics. For each analysis step, the incoming arrows indicate the sources of the input files, and the outgoing arrows indicate the next analysis step(s) to which the output files are passed. Dotted arrows indicate optional connections. The numbers in the legend indicate Cactus’ three main parts: 1. Preprocessing, 2. Differential Analysis, and 3. Enrichment Analysis. Abbreviations: ATAC, ATAC-Seq; mRNA, mRNA-Seq; DA, differential analysis; QC, quality controls; DASs, differential analysis subsets; ET, experiment type; TV, threshold value; FC, fold change; DAR, differentially accessible regions; PA, DAR peak annotation; KEGG, Kyoto Encyclopedia of Genes and Genomes; and GO, gene ontology.

Figure S5. Diagrams showing how differential analysis results are split by experiment type and fold change filters. Panel (a) shows the color code used in all panels, with blue circles representing genomic regions (either DARs or promoters of DEGs) and black circles representing gene sets (either the closest genes of DARs or DEGs). Enrichment of internal GRs and GSs indicates enrichment of GRs and GSs (i.e., DASs) in other GRs and GSs generated by the pipeline. Panels (b-g) show all possible splits of differential analysis results by experiment type (ATAC-Seq – turquoise, mRNA-Seq – orange, or both ATAC-Seq and mRNA-Seq – purple) and by fold change type (up – yellow, or down – green), with either an increase (b) or a decrease (c) in chromatin accessibility, an increase (d) or a decrease (e) in gene expression, and an increase (f) or a decrease (g) in both chromatin accessibility and gene expression. The HA-HE and LA-LE terminology has been previously described in [1]. Black lines and blue circles represent DNA and nucleosomes, respectively. Orange lines represent mRNA molecules.

Figure S6. Example of ATAC-Seq signal for a LA-LE gene (nhr-76, top panel) and a HA-HE gene (tsp-1, bottom panel). Purple and yellow tracks show the HGM-4 RNAi and the control condition, respectively. ATAC-Seq peaks are shown in the blue track, just above the gene model track, with exons shown in red. Please observe that it is not possible to show the signal for mRNA-Seq data since an alignment-free method is used for the quantification of gene expression levels.

### Supplementary tables

**Cactus results for reprogramming-promoting genes as defined in Figure 6L of the Kolundzic *et* *al.* study** [2]**.**

| **COMP** | **gene_name** | **gene_id** | **pval** | **padj** | **L2FC** | **FC** |
| --- | --- | --- | --- | --- | --- | --- |
| supt16h_vs_ctl | LIN28B | ENSG00000187772 | 0.046 | 0.12 | 1.35 | up |
| ssrp1_vs_ctl | BMP2 | ENSG00000125845 | 0.00004 | 0.0012 | 1.17 | up |
| supt16h_vs_ctl | BMP2 | ENSG00000125845 | 0.0004 | 0.0038 | 1.16 | up |
| ssrp1_vs_ctl | SALL4 | ENSG00000101115 | 0.0055 | 0.046 | 0.88 | up |
| ssrp1_vs_ctl | CEBPB | ENSG00000172216 | 0.000093 | 0.0022 | 0.82 | up |
| supt16h_vs_ctl | SALL4 | ENSG00000101115 | 0.01 | 0.041 | 0.78 | up |
| ssrp1_vs_ctl | FGF2 | ENSG00000138685 | 0.00023 | 0.0045 | 0.7 | up |
| supt16h_vs_ctl | KDM4C | ENSG00000107077 | 0.0055 | 0.027 | 0.47 | up |
| ssrp1_vs_ctl | KDM4C | ENSG00000107077 | 0.016 | 0.091 | 0.47 | up |
| supt16h_vs_ctl | PRDM15 | ENSG00000141956 | 0.05 | 0.13 | 0.47 | up |
| ssrp1_vs_ctl | LIN28B | ENSG00000187772 | 0.73 | 0.85 | 0.29 | up |
| ssrp1_vs_ctl | PRDM15 | ENSG00000141956 | 0.85 | 0.92 | 0.08 | up |
| supt16h_vs_ctl | FGF2 | ENSG00000138685 | 0.033 | 0.097 | -0.33 | down |
| supt16h_vs_ctl | CEBPB | ENSG00000172216 | 0.0042 | 0.022 | -0.66 | down |

**Cactus results for reprogramming-preventing genes as defined in Figure 6L of the Kolundzic *et* *al.* study** [2]**.**

| **COMP** | **gene_name** | **gene_id** | **pval** | **padj** | **L2FC** | **FC** |
| --- | --- | --- | --- | --- | --- | --- |
| supt16h_vs_ctl | PTPN11 | ENSG00000179295 | 0.056 | 0.14 | 0.31 | up |
| ssrp1_vs_ctl | DRAM1 | ENSG00000136048 | 0.9 | 0.95 | -0.02 | down |
| supt16h_vs_ctl | RBBP7 | ENSG00000102054 | 0.17 | 0.31 | -0.29 | down |
| supt16h_vs_ctl | DRAM1 | ENSG00000136048 | 0.068 | 0.16 | -0.31 | down |
| supt16h_vs_ctl | CHAF1B | ENSG00000159259 | 0.057 | 0.14 | -0.33 | down |
| ssrp1_vs_ctl | PTPN11 | ENSG00000179295 | 0.014 | 0.082 | -0.49 | down |
| supt16h_vs_ctl | SUV39H2 | ENSG00000152455 | 0.03 | 0.09 | -0.51 | down |
| supt16h_vs_ctl | SUV39H1 | ENSG00000101945 | 0.0029 | 0.017 | -0.58 | down |
| ssrp1_vs_ctl | CHAF1B | ENSG00000159259 | 0.0022 | 0.025 | -0.62 | down |
| ssrp1_vs_ctl | RBBP7 | ENSG00000102054 | 0.0005 | 0.008 | -0.77 | down |
| supt16h_vs_ctl | SUMO2 | ENSG00000188612 | 0.00000073 | 0.000034 | -0.78 | down |
| ssrp1_vs_ctl | NR2F1 | ENSG00000175745 | 0.008 | 0.059 | -0.88 | down |
| supt16h_vs_ctl | NR2F1 | ENSG00000175745 | 0.014 | 0.052 | -1.09 | down |
| ssrp1_vs_ctl | SUMO2 | ENSG00000188612 | 2.6E-11 | 4.2E-09 | -1.25 | down |
| ssrp1_vs_ctl | SUV39H2 | ENSG00000152455 | 0.0000004 | 0.000023 | -1.35 | down |
| ssrp1_vs_ctl | SUV39H1 | ENSG00000101945 | 0.00000006 | 0.0000046 | -1.77 | down |
| supt16h_vs_ctl | PTPN11 | ENSG00000179295 | 0.056 | 0.14 | 0.31 | up |
| ssrp1_vs_ctl | DRAM1 | ENSG00000136048 | 0.9 | 0.95 | -0.02 | Down |

Table S1. Gene expression results for reprogramming-promoting and -inhibitory genes. These tables show Cactus mRNA-Seq results for genes promoting (upper table) or preventing (lower table) reprogramming, as defined in Figure 6L of the *Kolundzic et* *al.* study. One reprogramming-promoting gene, ESRRB, was not detected by Cactus in the mRNA-Seq data and therefore is missing. KDM4C is an alias for the gene JMJD2C. Tables are sorted by descending Log_2_ Fold Change. Abbreviations: COMP, comparison; pval, p-value; padj, adjusted p-value (false discovery rate); L2FC, Log_2_ fold change; FC, direction of fold change; up, up-regulated gene; down, down-regulated gene; ctl, control.
