## Supplementary figures and images for "Cactus: a user-friendly and reproducible ATAC-Seq and mRNA-Seq analysis pipeline for data preprocessing, differential analysis, and enrichment analysis"

### Figure S1

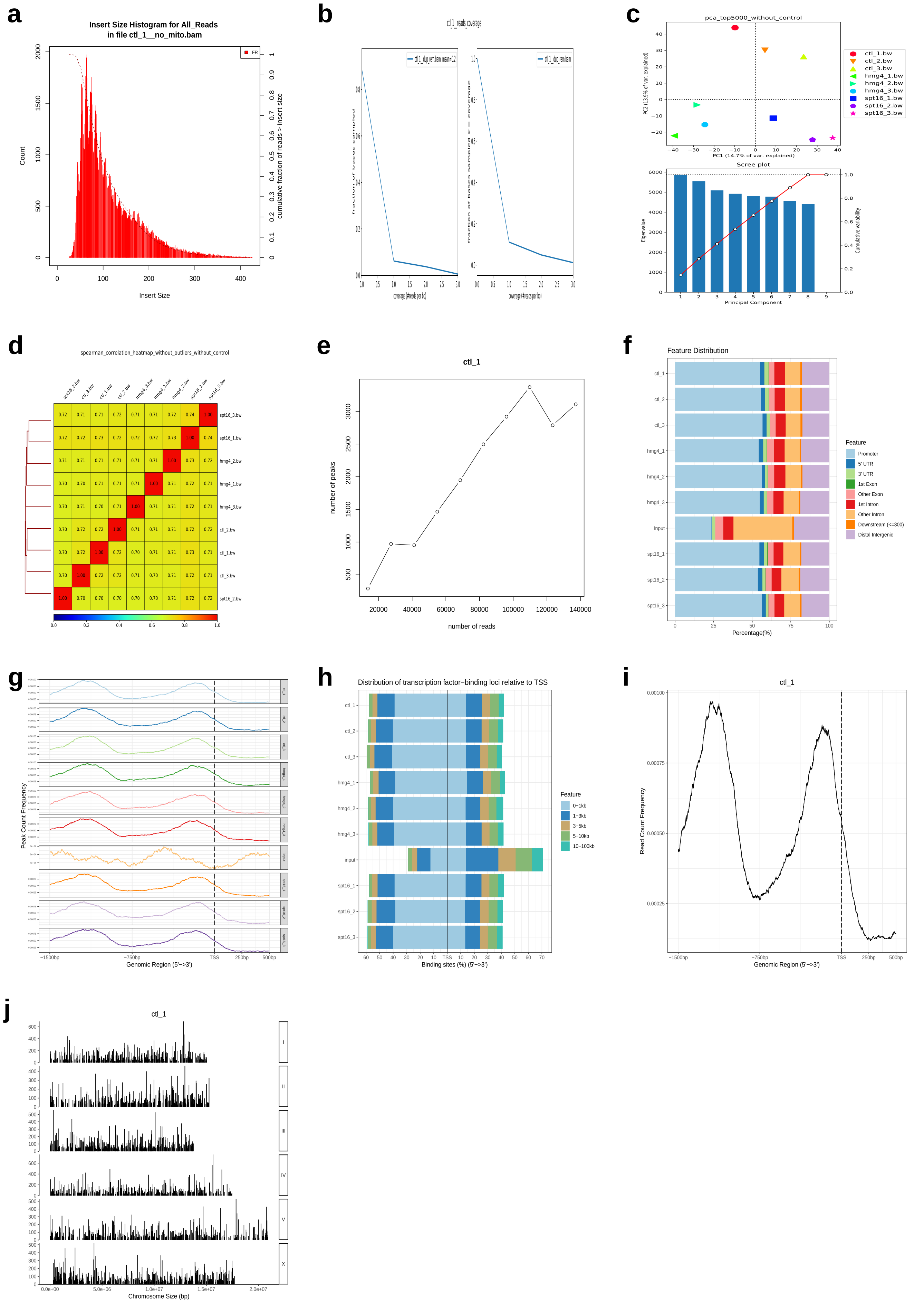

### Figure S4

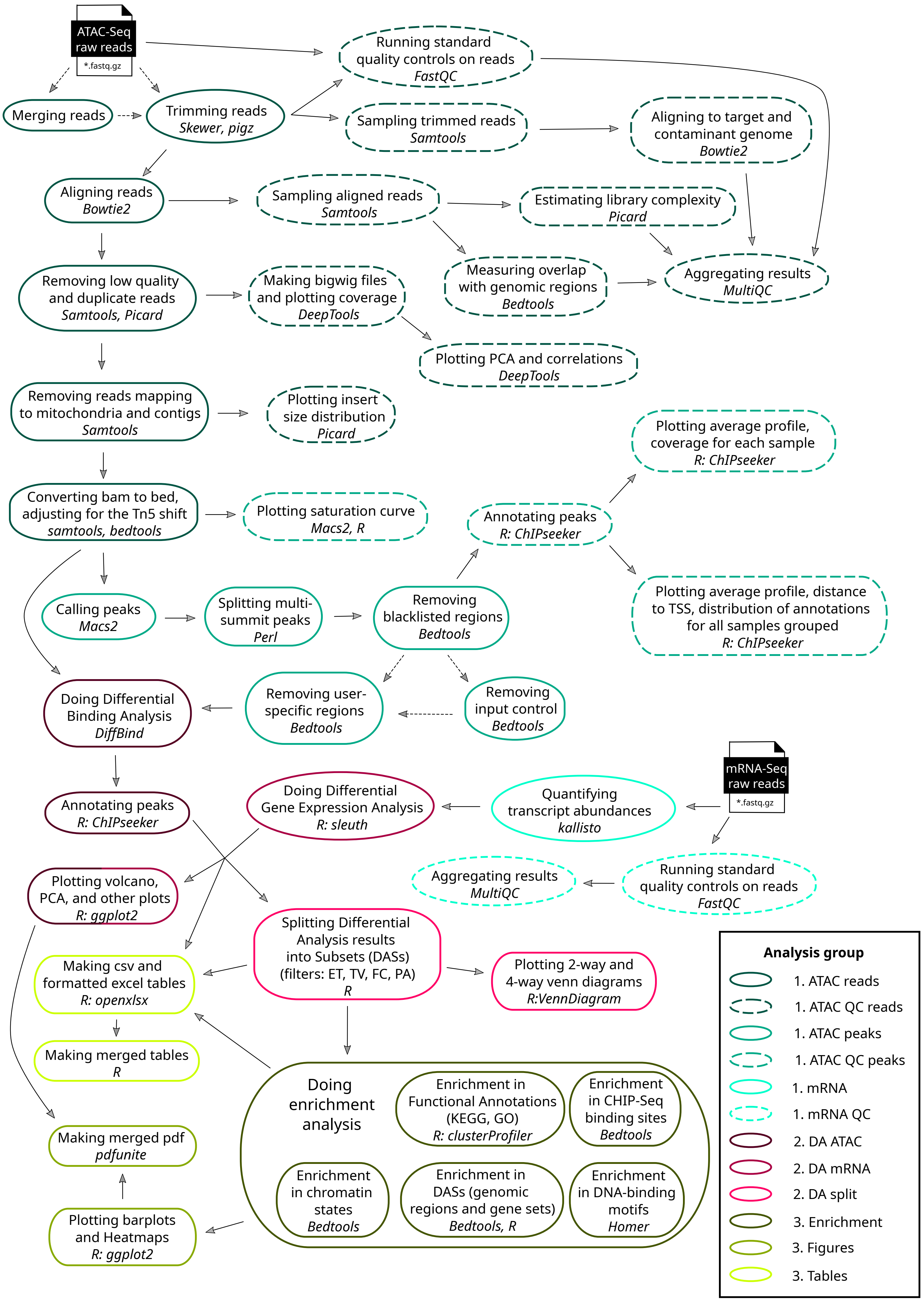

### Figure S5

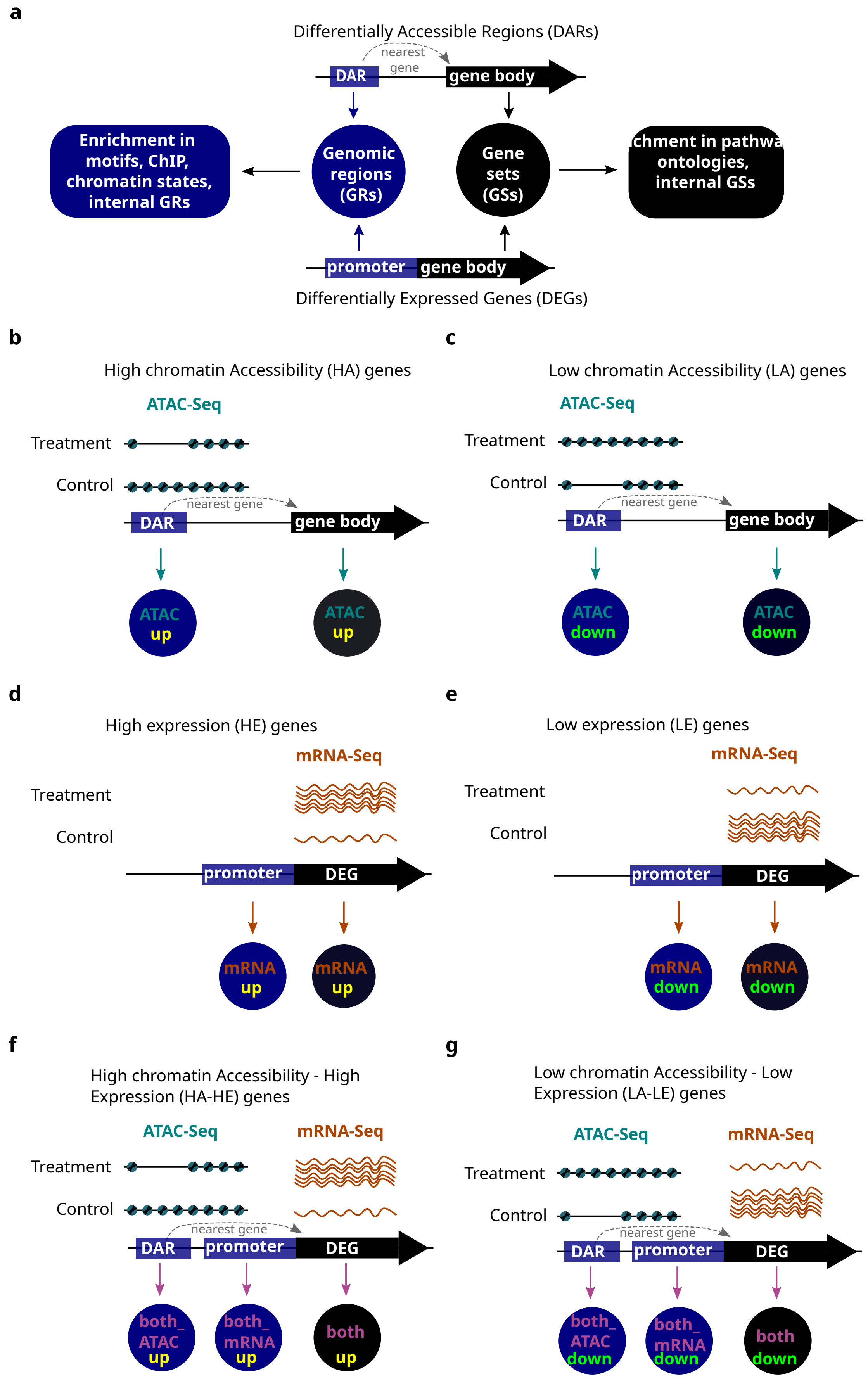

### Figure S6

#### Example of a LA−LE gene

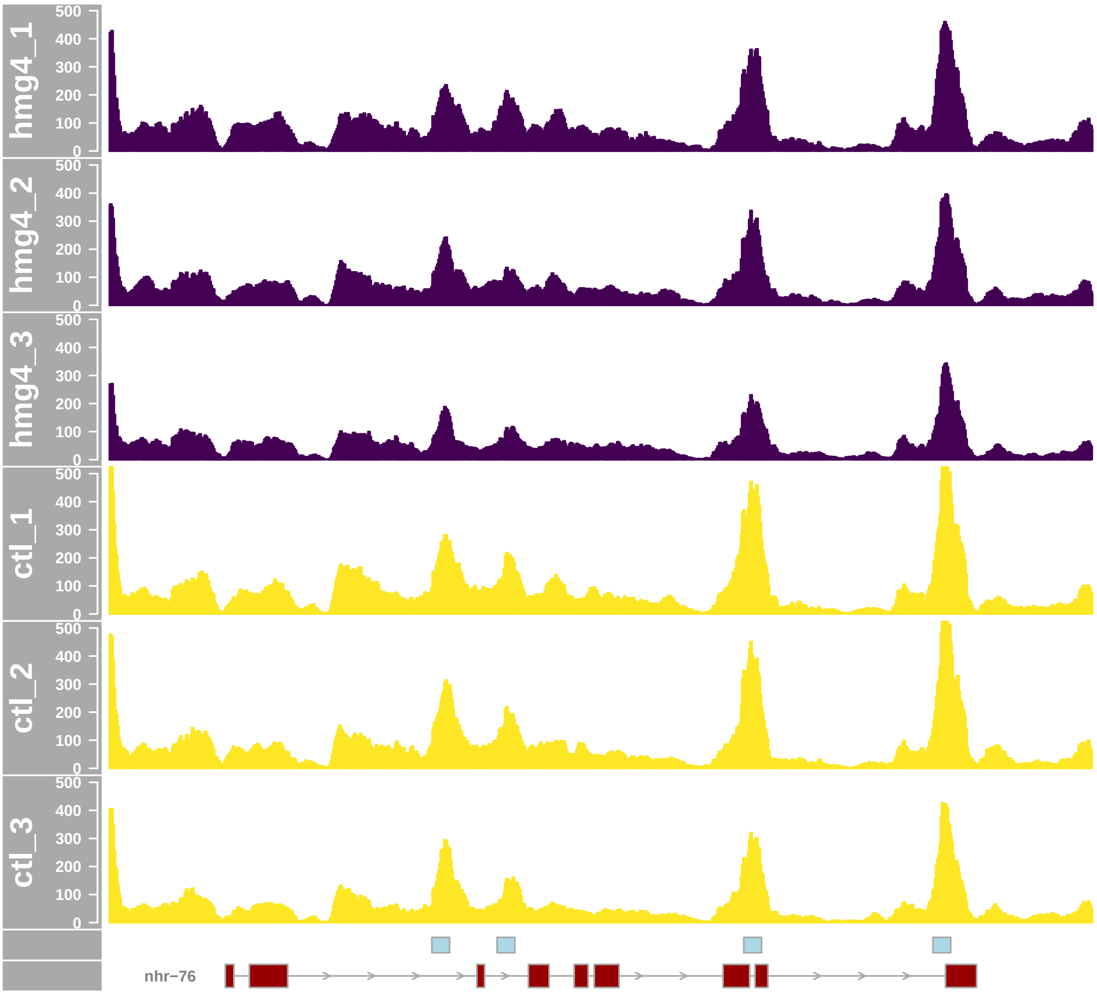


#### Example of a HA−HE gene

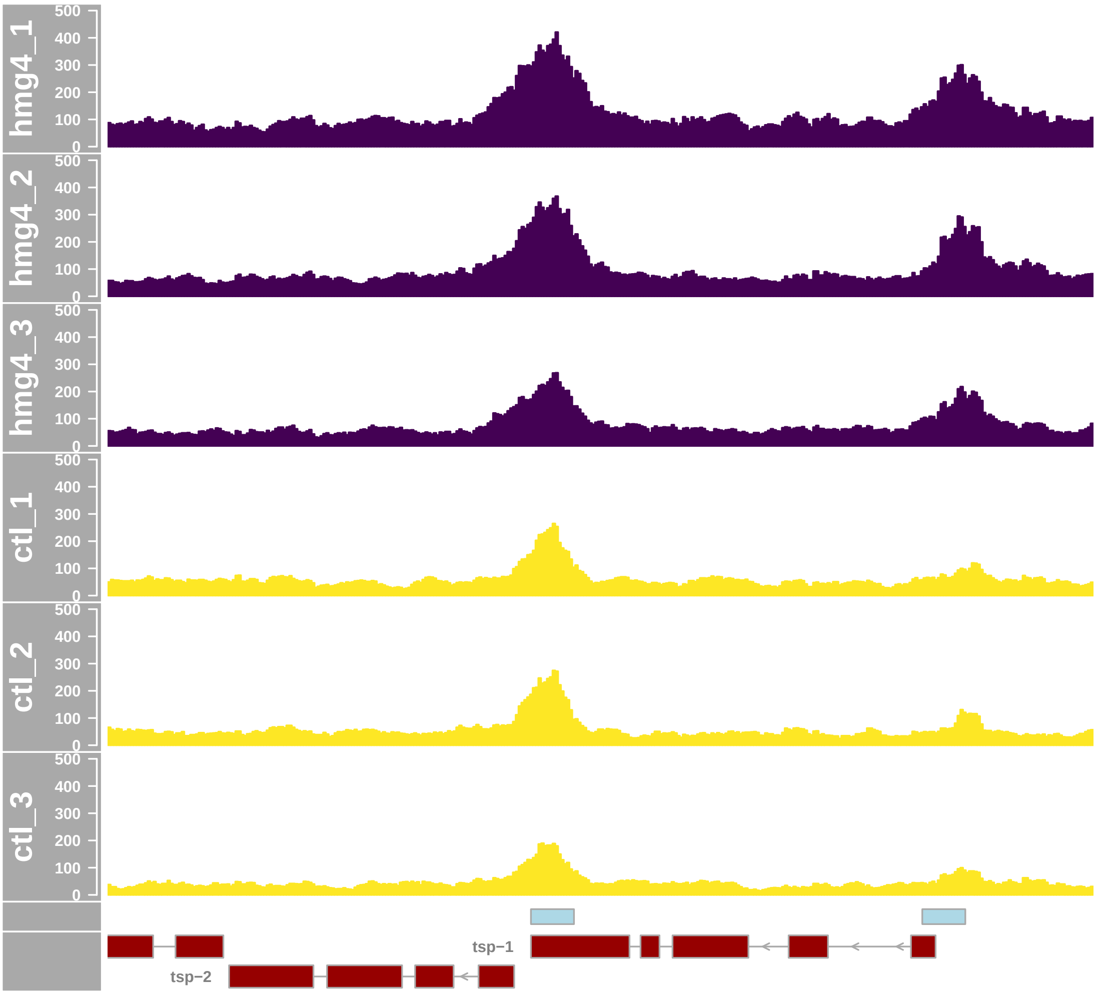
