## Supplementary material for "Cactus: a user-friendly and reproducible ATAC-Seq and mRNA-Seq analysis pipeline for data preprocessing, differential analysis, and enrichment analysis": Figure S2

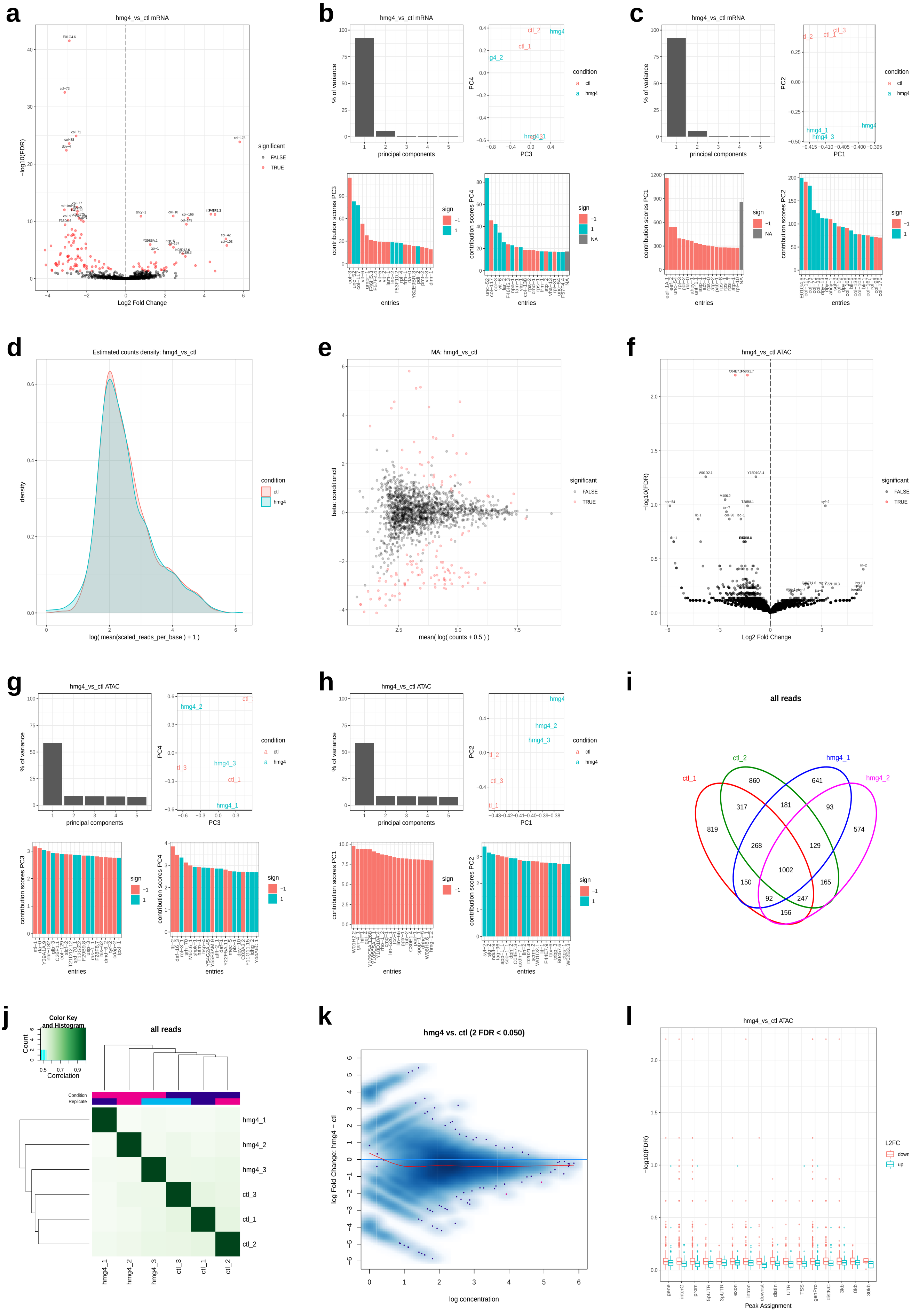
a
hmg4_vs_ctl mRNA
−log10(FDR)
Log2 Fold Change
significant
FALSE
TRUE
b
hmg4_vs_ctl mRNA
% of variance
principal components
PC4
PC3
condition
ctl
hmg4
contribution scores PC3
contribution scores PC4
entries
sign
c
hmg4_vs_ctl mRNA
PC2
PC1
contribution scores PC1
contribution scores PC2
d
Estimated counts density: hmg4_vs_ctl
density
log( mean(scaled_reads_per_base ) + 1 )
e
MA: hmg4_vs_ctl
beta: conditionctl
mean( log( counts + 0.5 ) )
f
hmg4_vs_ctl ATAC
g
hmg4_vs_ctl ATAC
h
hmg4_vs_ctl ATAC
i
all reads
ctl_1
ctl_2
hmg4_1
hmg4_2
j
Color Key and Histogram
Count
Correlation
all reads
Condition
Replicate
hmg4_1
hmg4_2
hmg4_3
ctl_3
ctl_1
ctl_2
k
hmg4 vs. ctl (2 FDR < 0.050)
log Fold Change: hmg4 − ctl
log concentration
l
hmg4_vs_ctl ATAC
L2FC
down
up
Peak Assignment

**m**

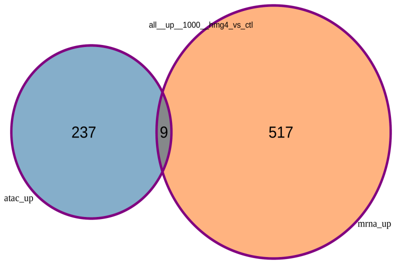


**n**

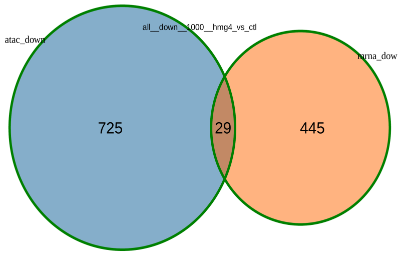


**o**

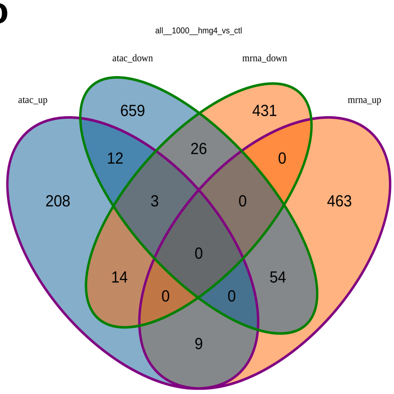


**p**

| | A | B | C | D | E | F | G | H | I | J | K | L | M |
|---|---|---|---|---|---|---|---|---|---|---|---|---|---|
| 1 | COMP | peak_id | chr | start | end | gene_name | gene_id | pval | padj | L2FC | distance_to_tss | annotation | conc |
| 2 | hmg4_vs_ctl | 5368 | X | 348223 | 348373 | C04E7.3 | WBGene00015430 | 1.9E-06 | 0.0063 | -2.04 | -1006 | Promoter | 3.92007 |
| 3 | hmg4_vs_ctl | 1854 | II | 12944962 | 12945112 | F58G1.7 | WBGene00010269 | 1.3E-06 | 0.0063 | -1.34 | 3146 | ntron (F58G1.7.1/WBGene00010269, intron 5 of 6 | 4.93086 |
| 4 | spt16_vs_ctl | 3487 | IV | 8260945 | 8261095 | vit-6 | WBGene00006930 | 2.1E-06 | 0.014 | -3.98 | -62 | Promoter | 2.77189 |
| 5 | hmg4_vs_spt16 | 659 | I | 10387293 | 10387443 | mgl-2 | WBGene00003233 | 1.2E-05 | 0.036 | -4.54 | 3945 | xon (F45H11.4c.1/WBGene00003233, exon 9 of 17 | 2.32765 |
| 6 | hmg4_vs_spt16 | 2174 | III | 7162007 | 7162157 | syf-2 | WBGene00019402 | 1.2E-05 | 0.036 | 4.44 | 0 | Promoter | 2.45464 |
| 7 | spt16_vs_ctl | 757 | I | 10387293 | 10387443 | mgl-2 | WBGene00003233 | 1.5E-05 | 0.047 | 4.5 | 3945 | xon (F45H11.4c.1/WBGene00003233, exon 9 of 17 | 2.32759 |
| 8 | spt16_vs_ctl | 756 | I | 10386831 | 10386981 | mgl-2 | WBGene00003233 | 2.4E-05 | 0.052 | 6.01 | 4407 | (on (F45H11.4c.1/WBGene00003233, exon 10 of 1 | 1.94437 |
| 9 | hmg4_vs_ctl | 1983 | II | 14815014 | 14815164 | W01D2.1 | WBGene00012179 | 3.3E-05 | 0.0551 | -3.77 | -354 | Promoter | 2.45952 |
| 10 | hmg4_vs_ctl | 901 | I | 12815911 | 12816061 | Y18D10A.4 | WBGene00012477 | 2.7E-05 | 0.0551 | -0.85 | 7 | Promoter | 5.56956 |

**q**

| | N | O | P | Q | R | S | T | U | V | W | X |
|---|---|---|---|---|---|---|---|---|---|---|---|
| 1 | conc_cond1 | conc_cond2 | counts_cond1 | counts_cond2 | jbrowse | gene_chr | gene_start | gene_end | gene_width | gene_strand | entrez_id |
| 2 | 2.5843 | 4.60158 | 7.74\|2.97\|6.96 | 27.29\|34.2\|12.41 | X:348,223..348,373 | X | 334567 | 347217 | 12650 | - | 259695 |
| 3 | 4.14316 | 5.4375 | 16.45\|14.85\|20.87 | 46.5\|41.25\|44.47 | II:12,944,962..12,945,112 | II | 12943849 | 12948258 | 4409 | - | 174937 |
| 4 | 0 | 3.69994 | 1.97\|0\|0 | 12.55\|13.45\|11.85 | IV:8,260,945..8,261,095 | IV | 8261157 | 8266459 | 5302 | + | 177619 |
| 5 | 0 | 3.27892 | 0\|0.96\|0 | 8.03\|8.5\|14.09 | I:10,387,293..10,387,443 | I | 10383144 | 10391388 | 8244 | - | 172914 |
| 6 | 3.41007 | 0 | 16.92\|6.73\|6.76 | 0\|1.06\|0 | III:7,162,007..7,162,157 | III | 7161160 | 7162115 | 955 | - | 176003 |
| 7 | 3.2789 | 0 | 7.9\|8.35\|13.85 | 0.97\|0\|0 | I:10,387,293..10,387,443 | I | 10383144 | 10391388 | 8244 | - | 172914 |
| 8 | 2.94437 | 0 | 5.92\|8.35\|9.59 | 0\|0\|0 | I:10,386,831..10,386,981 | I | 10383144 | 10391388 | 8244 | - | 172914 |
| 9 | 0 | 3.36946 | 1.94\|0\|0 | 13.14\|8.05\|10.34 | II:14,815,014..14,815,164 | II | 14815518 | 14816140 | 622 | + | 175141 |
| 10 | 5.11495 | 5.91472 | 47.41\|33.65\|20.87 | 71.77\|57.34\|54.81 | I:12,815,911..12,816,061 | I | 12815904 | 12816558 | 654 | + | 189473 |

**r**

| | X | Y | Z | AA | AB | AC | AD | AE | AF | AG | AH | AI | AJ | AK | AL | AM | AN | AO | AP |
|---|---|---|---|---|---|---|---|---|---|---|---|---|---|---|---|---|---|---|---|
| 1 | entrez_id | FC_up | FC_down | PA_gene | PA_interG | PA_prom | PA_5pUTR | PA_3pUTR | PA_exon | PA_intron | PA_downst | PA_distIn | PA_UTR | PA_TSS | PA_genPro | PA_distNC | PA_3kb | PA_8kb | PA_30kb |
| 2 | 259695 | FALSE | TRUE | FALSE | TRUE | TRUE | FALSE | FALSE | FALSE | FALSE | FALSE | FALSE | FALSE | FALSE | TRUE | FALSE | FALSE | FALSE | FALSE |
| 3 | 174937 | FALSE | TRUE | TRUE | FALSE | FALSE | FALSE | FALSE | FALSE | TRUE | FALSE | FALSE | FALSE | FALSE | TRUE | TRUE | TRUE | FALSE | FALSE |
| 4 | 177619 | FALSE | TRUE | FALSE | TRUE | TRUE | FALSE | FALSE | FALSE | FALSE | FALSE | FALSE | FALSE | FALSE | TRUE | FALSE | FALSE | FALSE | FALSE |
| 5 | 172914 | FALSE | TRUE | TRUE | FALSE | FALSE | FALSE | FALSE | TRUE | FALSE | FALSE | FALSE | FALSE | FALSE | TRUE | FALSE | TRUE | FALSE | FALSE |
| 6 | 176003 | TRUE | FALSE | TRUE | FALSE | TRUE | FALSE | FALSE | TRUE | FALSE | FALSE | FALSE | FALSE | TRUE | TRUE | FALSE | FALSE | FALSE | FALSE |
| 7 | 172914 | TRUE | FALSE | TRUE | FALSE | FALSE | FALSE | FALSE | TRUE | FALSE | FALSE | FALSE | FALSE | FALSE | TRUE | FALSE | TRUE | FALSE | FALSE |
| 8 | 172914 | TRUE | FALSE | TRUE | FALSE | FALSE | FALSE | FALSE | TRUE | TRUE | FALSE | FALSE | FALSE | FALSE | TRUE | FALSE | TRUE | FALSE | FALSE |
| 9 | 175141 | FALSE | TRUE | FALSE | TRUE | TRUE | FALSE | FALSE | FALSE | FALSE | TRUE | FALSE | FALSE | FALSE | TRUE | FALSE | FALSE | FALSE | FALSE |
| 10 | 189473 | FALSE | TRUE | TRUE | FALSE | TRUE | FALSE | FALSE | TRUE | FALSE | FALSE | FALSE | FALSE | FALSE | TRUE | FALSE | FALSE | FALSE | FALSE |

**s**

| | A | B | C | D | E | F | G | H | I | J | K | L |
|---|---|---|---|---|---|---|---|---|---|---|---|---|
| 1 | COMP | chr | start | end | width | strand | gene_name | gene_id | entrez_id | pval | padj | L2FC |
| 2 | hmg4_vs_ctl | II | 13482405 | 13485780 | 3375 | + | E01G4.6 | WBGene00008448 | 174990 | 1.8E-45 | 3E-42 | -2.89 |
| 3 | hmg4_vs_ctl | II | 4872098 | 4873129 | 1031 | - | col-73 | WBGene00000649 | 184366 | 3.6E-36 | 3E-33 | -3.12 |
| 4 | spt16_vs_ctl | V | 9164292 | 9165307 | 1015 | - | col-10 | WBGene00000599 | 179298 | 1.6E-34 | 2.7E-31 | 3.65 |
| 5 | hmg4_vs_spt16 | V | 9629943 | 9631012 | 1069 | + | col-146 | WBGene00000719 | 188178 | 8E-34 | 1.3E-30 | -5.79 |
| 6 | spt16_vs_ctl | II | 12172573 | 12176069 | 3496 | + | rol-1 | WBGene00004394 | 174857 | 8.8E-33 | 7.4E-30 | -6.54 |
| 7 | hmg4_vs_ctl | II | 3500077 | 3502570 | 2493 | - | col-71 | WBGene00000647 | 173695 | 2.2E-28 | 1.2E-25 | -2.54 |
| 8 | hmg4_vs_ctl | X | 10077597 | 10078728 | 1131 | + | col-176 | WBGene00000749 | 181219 | 3.2E-27 | 1.3E-24 | 5.81 |
| 9 | hmg4_vs_ctl | II | 8568035 | 8569088 | 1053 | - | col-38 | WBGene00000615 | 174370 | 7.8E-27 | 2.6E-24 | -2.89 |
| 10 | hmg4_vs_ctl | IV | 14985543 | 14986784 | 1241 | - | dpy-4 | WBGene00001066 | 178414 | 1.4E-25 | 3.9E-23 | -3.04 |
| 11 | spt16_vs_ctl | II | 3500077 | 3502570 | 2493 | - | col-71 | WBGene00000647 | 173695 | 8.9E-24 | 4.5E-21 | -5.56 |
| 12 | spt16_vs_ctl | V | 12352745 | 12354320 | 1575 | + | sqt-3 | WBGene00005018 | 179693 | 1.1E-23 | 4.5E-21 | -3.14 |
| 13 | spt16_vs_ctl | II | 5651394 | 5652434 | 1040 | - | bli-2 | WBGene00000252 | 191611 | 1.5E-22 | 5.1E-20 | -5.6 |
| 14 | spt16_vs_ctl | X | 6280016 | 6280921 | 905 | + | col-166 | WBGene00000739 | 180902 | 8.5E-22 | 2.4E-19 | 4.01 |
| 15 | spt16_vs_ctl | IV | 14985543 | 14986784 | 1241 | - | dpy-4 | WBGene00001066 | 178414 | 4.8E-21 | 1.2E-18 | -1.93 |

**t**

| | M | N | O | P | Q | R | S | T |
|---|---|---|---|---|---|---|---|---|
| 1 | TXCHROM | se_b | mean_obs | var_obs | tech_var | sigma_sq | smooth_sigma_sq | final_sigma_sq |
| 2 | II | 0.204 | 6.122 | 2.53208 | 0.04643 | -0.007429 | 0.01601 | 0.01601 |
| 3 | II | 0.2487 | 5.843 | 2.98578 | 0.07296 | 0.00308 | 0.01979 | 0.01979 |
| 4 | V | 0.2977 | 5.982 | 4.03557 | 0.11799 | -0.063073 | 0.0149 | 0.0149 |
| 5 | V | 0.4773 | 1.893 | 10.0713 | 0.05409 | -0.018559 | 0.28761 | 0.28761 |
| 6 | II | 0.5486 | 2.801 | 13.201 | 0.20212 | 0.249338 | 0.11523 | 0.24934 |
| 7 | II | 0.2298 | 5.749 | 1.96469 | 0.05804 | -0.019021 | 0.02116 | 0.02116 |
| 8 | X | 0.5378 | 2.943 | 10.465 | 0.35832 | 0.060837 | 0.07546 | 0.07546 |
| 9 | II | 0.2696 | 5.43 | 2.57898 | 0.08298 | 0.005244 | 0.02607 | 0.02607 |
| 10 | IV | 0.2906 | 5.198 | 2.8167 | 0.09677 | -0.035671 | 0.02988 | 0.02988 |
| 11 | II | 0.5534 | 4.257 | 9.62506 | 0.41614 | 0.008258 | 0.04325 | 0.04325 |
| 12 | V | 0.3124 | 5.127 | 3.01891 | 0.12054 | -0.034378 | 0.02588 | 0.02588 |
| 13 | II | 0.5733 | 2.883 | 9.76933 | 0.38748 | 0.057785 | 0.10556 | 0.10556 |
| 14 | X | 0.4184 | 5.229 | 4.89153 | 0.23828 | -0.166589 | 0.02431 | 0.02431 |
| 15 | IV | 0.2055 | 5.769 | 1.14742 | 0.04615 | -0.015078 | 0.0172 | 0.0172 |

**u**

| | A | B | C | D | E | F | G | H | I | J | K | L |
|---|---|---|---|---|---|---|---|---|---|---|---|---|
| 1 | ET | PF | FC | TV | COMP | peak_id | chr | gene_name | gene_id | pval | padj | L2FC |
| 2 | mRNA | Null | down | 200 | hmg4_vs_ctl | Null | II | E01G4.6 | WBGene00008448 | 1.8E-45 | 3E-42 | -2.89 |
| 3 | mRNA | Null | down | 1000 | hmg4_vs_ctl | Null | II | E01G4.6 | WBGene00008448 | 1.8E-45 | 3E-42 | -2.89 |
| 4 | mRNA | Null | down | 200 | hmg4_vs_ctl | Null | II | col-73 | WBGene00000649 | 3.6E-36 | 3E-33 | -3.12 |
| 5 | mRNA | Null | down | 1000 | hmg4_vs_ctl | Null | II | col-73 | WBGene00000649 | 3.6E-36 | 3E-33 | -3.12 |
| 6 | mRNA | Null | up | 200 | spt16_vs_ctl | Null | V | col-10 | WBGene00000599 | 1.6E-34 | 2.7E-31 | 3.65 |
| 7 | mRNA | Null | up | 1000 | spt16_vs_ctl | Null | V | col-10 | WBGene00000599 | 1.6E-34 | 2.7E-31 | 3.65 |
| 8 | mRNA | Null | down | 200 | hmg4_vs_spt16 | Null | V | col-146 | WBGene00000719 | 8E-34 | 1.3E-30 | -5.79 |
| 9 | mRNA | Null | down | 1000 | hmg4_vs_spt16 | Null | V | col-146 | WBGene00000719 | 8E-34 | 1.3E-30 | -5.79 |
| 10 | mRNA | Null | down | 200 | spt16_vs_ctl | Null | II | rol-1 | WBGene00004394 | 8.8E-33 | 7.4E-30 | -6.54 |
| 11 | mRNA | Null | down | 1000 | spt16_vs_ctl | Null | II | rol-1 | WBGene00004394 | 8.8E-33 | 7.4E-30 | -6.54 |
| 12 | mRNA | Null | down | 200 | hmg4_vs_ctl | Null | II | col-71 | WBGene00000647 | 2.2E-28 | 1.2E-25 | -2.54 |
| 13 | mRNA | Null | down | 1000 | hmg4_vs_ctl | Null | II | col-71 | WBGene00000647 | 2.2E-28 | 1.2E-25 | -2.54 |
| 14 | mRNA | Null | up | 200 | hmg4_vs_ctl | Null | X | col-176 | WBGene00000749 | 3.2E-27 | 1.3E-24 | 5.81 |
| 15 | mRNA | Null | up | 1000 | hmg4_vs_ctl | Null | X | col-176 | WBGene00000749 | 3.2E-27 | 1.3E-24 | 5.81 |

**v**

| | A | B | C | D | E | F | G | H | I | J | K | L |
|---|---|---|---|---|---|---|---|---|---|---|---|---|
| 1 | ET | PF | FC | TV | COMP | peak_id | chr | gene_name | gene_id | pval | padj | L2FC |
| 2 | mRNA | Null | down | 200\|1000 | hmg4_vs_ctl | Null | II | E01G4.6 | WBGene00008448 | 1.8E-45 | 3E-42 | -2.89 |
| 3 | mRNA | Null | down | 200\|1000 | hmg4_vs_ctl | Null | II | col-73 | WBGene00000649 | 3.6E-36 | 3E-33 | -3.12 |
| 4 | mRNA | Null | up | 200\|1000 | spt16_vs_ctl | Null | V | col-10 | WBGene00000599 | 1.6E-34 | 2.7E-31 | 3.65 |
| 5 | mRNA | Null | down | 200\|1000 | hmg4_vs_spt16 | Null | V | col-146 | WBGene00000719 | 8E-34 | 1.3E-30 | -5.79 |
| 6 | mRNA | Null | down | 200\|1000 | spt16_vs_ctl | Null | II | rol-1 | WBGene00004394 | 8.8E-33 | 7.4E-30 | -6.54 |
| 7 | mRNA | Null | down | 200\|1000 | hmg4_vs_ctl | Null | II | col-71 | WBGene00000647 | 2.2E-28 | 1.2E-25 | -2.54 |
| 8 | mRNA | Null | up | 200\|1000 | hmg4_vs_ctl | Null | X | col-176 | WBGene00000749 | 3.2E-27 | 1.3E-24 | 5.81 |
| 9 | mRNA | Null | down | 200\|1000 | hmg4_vs_ctl | Null | II | col-38 | WBGene00000615 | 7.8E-27 | 2.6E-24 | -2.89 |
| 10 | mRNA | Null | down | 200\|1000 | hmg4_vs_ctl | Null | IV | dpy-4 | WBGene00001066 | 1.4E-25 | 3.9E-23 | -3.04 |
| 11 | mRNA | Null | down | 200\|1000 | spt16_vs_ctl | Null | II | col-71 | WBGene00000647 | 8.9E-24 | 4.5E-21 | -5.56 |
| 12 | mRNA | Null | down | 200\|1000 | spt16_vs_ctl | Null | V | sqt-3 | WBGene00005018 | 1.1E-23 | 4.5E-21 | -3.14 |
| 13 | mRNA | Null | down | 200\|1000 | spt16_vs_ctl | Null | II | bli-2 | WBGene00000252 | 1.5E-22 | 5.1E-20 | -5.6 |
| 14 | mRNA | Null | up | 200\|1000 | spt16_vs_ctl | Null | X | col-166 | WBGene00000739 | 8.5E-22 | 2.4E-19 | 4.01 |
| 15 | mRNA | Null | down | 200\|1000 | spt16_vs_ctl | Null | IV | dpy-4 | WBGene00001066 | 4.8E-21 | 1.2E-18 | -1.93 |
