## Supplementary material for "Cactus: a user-friendly and reproducible ATAC-Seq and mRNA-Seq analysis pipeline for data preprocessing, differential analysis, and enrichment analysis": Figure S3

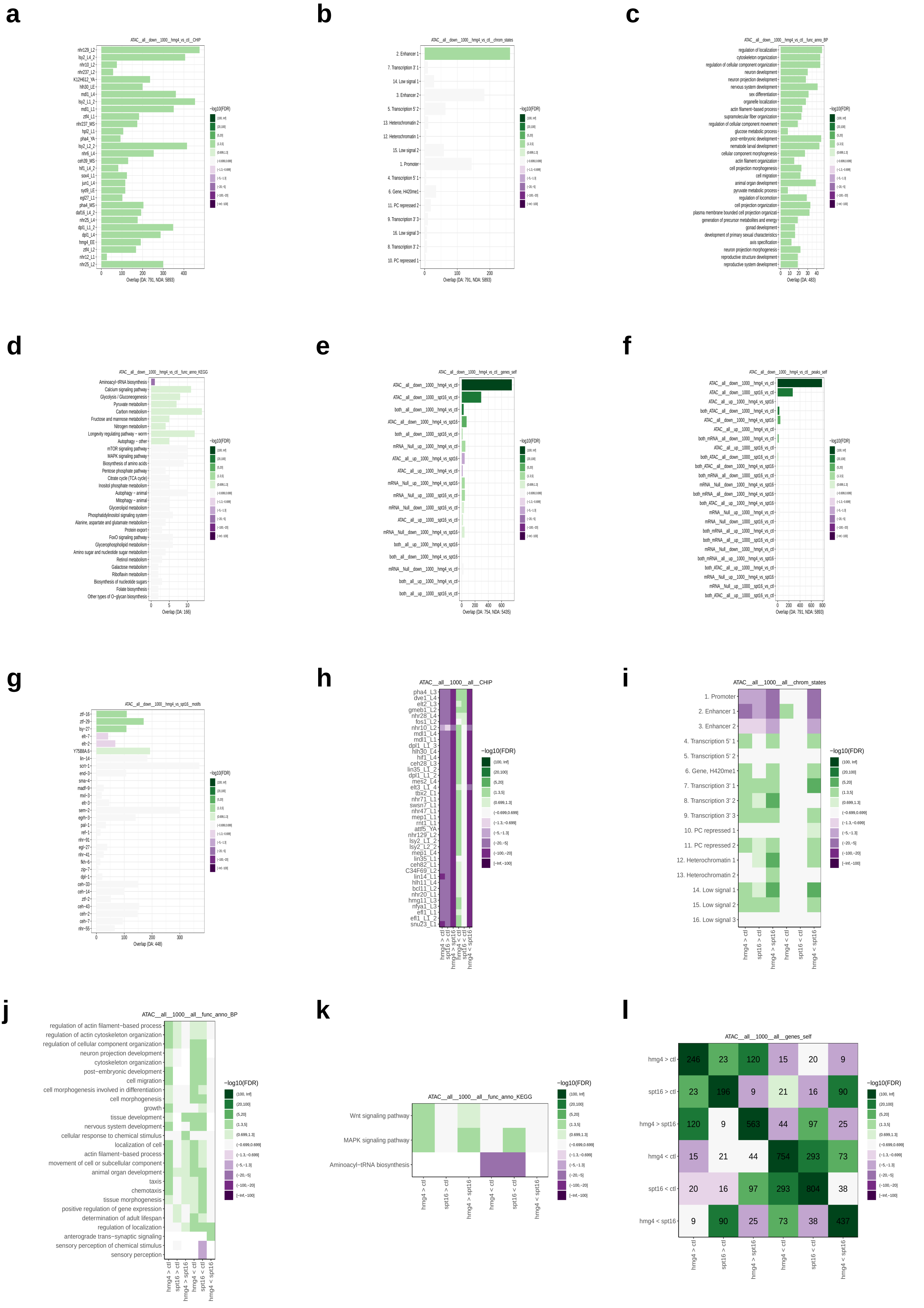
a
ATAC__all__down__1000__hmg4_vs_ctl__CHIP
Overlap (DA: 791, NDA: 5893)
b
ATAC__all__down__1000__hmg4_vs_ctl__chrom_states
2. Enhancer 1
7. Transcription 3' 1
14. Low signal 1
3. Enhancer 2
5. Transcription 5' 2
13. Heterochromatin 2
12. Heterochromatin 1
15. Low signal 2
1. Promoter
4. Transcription 5' 1
6. Gene, H420me1
11. PC repressed 2
9. Transcription 3' 3
16. Low signal 3
8. Transcription 3' 2
10. PC repressed 1
Overlap (DA: 791, NDA: 5893)
c
ATAC__all__down__1000__hmg4_vs_ctl__func_anno_BP
Overlap (DA: 483)
d
ATAC__all__down__1000__hmg4_vs_ctl__func_anno_KEGG
Overlap (DA: 166)
e
ATAC__all__down__1000__hmg4_vs_ctl__genes_self
Overlap (DA: 754, NDA: 5435)
f
ATAC__all__down__1000__hmg4_vs_ctl__peaks_self
Overlap (DA: 791, NDA: 5893)
g
ATAC__all__down__1000__hmg4_vs_spt16__motifs
Overlap (DA: 448)
h
ATAC__all__1000__all__CHIP
i
ATAC__all__1000__all__chrom_states
j
ATAC__all__1000__all__func_anno_BP
regulation of actin filament−based process
regulation of actin cytoskeleton organization
regulation of cellular component organization
neuron projection development
cytoskeleton organization
post−embryonic development
cell migration
cell morphogenesis involved in differentiation
cell morphogenesis
growth
tissue development
nervous system development
cellular response to chemical stimulus
localization of cell
actin filament−based process
movement of cell or subcellular component
animal organ development
taxis
chemotaxis
tissue morphogenesis
positive regulation of gene expression
determination of adult lifespan
regulation of localization
anterograde trans−synaptic signaling
sensory perception of chemical stimulus
sensory perception
k
ATAC__all__1000__all__func_anno_KEGG
Wnt signaling pathway
MAPK signaling pathway
Aminoacyl−tRNA biosynthesis
l
ATAC__all__1000__all__genes_self
hmg4 > ctl
spt16 > ctl
hmg4 > spt16
hmg4 < ctl
spt16 < ctl
hmg4 < spt16
246 23 120 15 20 9
23 196 9 21 16 90
120 9 563 44 97 25
15 21 44 754 293 73
20 16 97 293 804 38
9 90 25 73 38 437
−log10(FDR)
(100, Inf]
(20,100]
(5,20]
(1.3,5]
(0.699,1.3]
(−0.699,0.699]
(−1.3,−0.699]
(−5,−1.3]
(−20,−5]
(−100,−20]
[−Inf,−100]

m

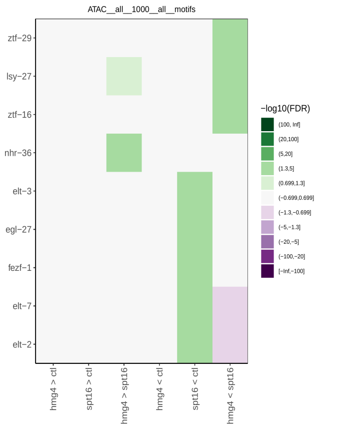


n

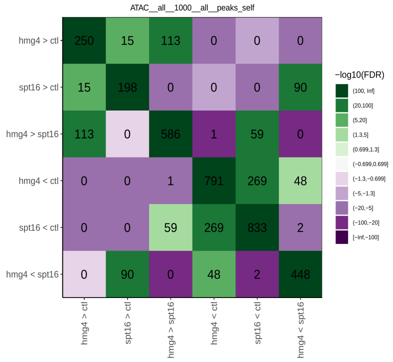


o

| | A | B | C | D | E | F | G | H | I | J | K | L | M | N | O | P |
|---|---|---|---|---|---|---|---|---|---|---|---|---|---|---|---|---|
| 1 | ET | PF | FC | TV | COMP | tgt | pval | padj | L2OR | pt_da | ov_da | tot_da | pt_nda | ov_nda | tot_nda | tot_tgt |
| 2 | ATAC | all | up | 1000 | hmg4_vs_spt16 | snu23_L1 | 5.7E-67 | 2.7E-64 | -2.36 | 22.15 | 128 | 578 | 59.36 | 3100 | 5222 | 6675 |
| 3 | ATAC | all | up | 1000 | hmg4_vs_spt16 | efl1_L1_3 | 5.6E-65 | 1.3E-62 | -2.57 | 14.01 | 81 | 578 | 49.18 | 2568 | 5222 | 4835 |
| 4 | ATAC | all | up | 1000 | hmg4_vs_spt16 | nfya1_L3 | 4E-63 | 6.3E-61 | -2.34 | 19.9 | 115 | 578 | 55.71 | 2909 | 5222 | 6932 |
| 5 | ATAC | all | up | 1000 | hmg4_vs_spt16 | efl1_L1 | 1E-62 | 1.2E-60 | -2.58 | 12.98 | 75 | 578 | 47.11 | 2460 | 5222 | 4534 |
| 6 | ATAC | all | up | 1000 | hmg4_vs_spt16 | nhr129_L2 | 3.9E-59 | 3.7E-57 | -2.2 | 23.36 | 135 | 578 | 58.35 | 3047 | 5222 | 6626 |
| 7 | ATAC | all | up | 1000 | hmg4_vs_spt16 | mep1_L4 | 1.6E-58 | 1.3E-56 | -2.39 | 15.57 | 90 | 578 | 49.08 | 2563 | 5222 | 5229 |
| 8 | ATAC | all | up | 1000 | hmg4_vs_spt16 | lin14_L1 | 9.4E-58 | 6.4E-56 | -2.11 | 27.68 | 160 | 578 | 62.37 | 3257 | 5222 | 11507 |
| 9 | ATAC | all | up | 1000 | hmg4_vs_spt16 | mes2_L4 | 2.4E-56 | 1.4E-54 | -2.38 | 14.71 | 85 | 578 | 47.3 | 2470 | 5222 | 5501 |
| 10 | ATAC | all | up | 1000 | hmg4_vs_spt16 | lin35_L1 | 3.1E-56 | 1.6E-54 | -2.17 | 21.97 | 127 | 578 | 55.88 | 2918 | 5222 | 6712 |
| 11 | ATAC | all | up | 1000 | hmg4_vs_spt16 | hlh30_L4 | 6.6E-56 | 3.1E-54 | -2.13 | 23.7 | 137 | 578 | 57.68 | 3012 | 5222 | 7168 |
| 12 | ATAC | all | up | 1000 | hmg4_vs_spt16 | C34F69_L2 | 3.5E-55 | 1.5E-53 | -2.23 | 18.69 | 108 | 578 | 51.82 | 2706 | 5222 | 5692 |
| 13 | ATAC | all | up | 1000 | hmg4_vs_spt16 | hmg11_L3 | 1.8E-54 | 7.3E-53 | -2.4 | 13.32 | 77 | 578 | 44.83 | 2341 | 5222 | 4037 |
| 14 | ATAC | all | up | 1000 | hmg4_vs_spt16 | ceh82_L1_2 | 1.4E-53 | 5.1E-52 | -2.44 | 12.28 | 71 | 578 | 43.18 | 2255 | 5222 | 3913 |
| 15 | ATAC | all | up | 1000 | hmg4_vs_spt16 | bcl11_L2 | 1.8E-53 | 5.9E-52 | -2.23 | 17.3 | 100 | 578 | 49.58 | 2589 | 5222 | 5133 |

p

| | A | B | C | D | E | F | G | H | I | J | K | L | M | N | O | P |
|---|---|---|---|---|---|---|---|---|---|---|---|---|---|---|---|---|
| 1 | ET | PF | FC | TV | COMP | tgt | pval | padj | L2OR | pt_da | ov_da | tot_da | pt_nda | ov_nda | tot_nda | tot_tgt |
| 5 | ATAC | all | up | 1000 | hmg4_vs_spt16 | 2. Enhancer 1 | 2.8E-18 | 4.5E-17 | -1.42 | 14.01 | 81 | 578 | 30.35 | 1585 | 5222 | 2785 |
| 12 | ATAC | all | up | 1000 | hmg4_vs_spt16 | 1. Promoter | 5.5E-14 | 4.4E-13 | -1.48 | 8.48 | 49 | 578 | 20.53 | 1072 | 5222 | 2661 |
| 14 | ATAC | all | down | 1000 | hmg4_vs_spt16 | 7. Transcription 3' 1 | 2.2E-13 | 3.5E-12 | 2.35 | 8.53 | 39 | 457 | 1.8 | 96 | 5343 | 3424 |
| 15 | ATAC | all | down | 1000 | hmg4_vs_spt16 | 2. Enhancer 1 | 7.1E-13 | 5.7E-12 | -1.29 | 14.88 | 68 | 457 | 29.91 | 1598 | 5343 | 2785 |
| 20 | ATAC | all | up | 1000 | hmg4_vs_ctl | 2. Enhancer 1 | 9.9E-11 | 1.6E-09 | -1.71 | 10.79 | 26 | 241 | 28.34 | 1826 | 6443 | 2785 |
| 21 | ATAC | all | up | 200 | hmg4_vs_spt16 | 2. Enhancer 1 | 1.2E-10 | 1.9E-09 | -2.77 | 5.69 | 7 | 123 | 29.22 | 1659 | 5677 | 2785 |
| 26 | ATAC | all | up | 1000 | hmg4_vs_spt16 | 14. Low signal 1 | 6.5E-10 | 3.5E-09 | 1.5 | 10.38 | 60 | 578 | 3.94 | 206 | 5222 | 1880 |
| 32 | ATAC | all | up | 1000 | hmg4_vs_spt16 | 12. Heterochromatin 1 | 4E-08 | 1.6E-07 | 2.54 | 3.46 | 20 | 578 | 0.61 | 32 | 5222 | 1117 |
| 34 | ATAC | all | up | 1000 | hmg4_vs_spt16 | 8. Transcription 3' 2 | 8.4E-08 | 2.7E-07 | 2.45 | 3.46 | 20 | 578 | 0.65 | 34 | 5222 | 1238 |
| 38 | ATAC | all | down | 1000 | hmg4_vs_spt16 | 1. Promoter | 3.5E-07 | 1.9E-06 | -1.06 | 10.72 | 49 | 457 | 20.06 | 1072 | 5343 | 2661 |
| 45 | ATAC | all | down | 1000 | hmg4_vs_spt16 | 14. Low signal 1 | 1.8E-06 | 7.1E-06 | 1.3 | 9.63 | 44 | 457 | 4.15 | 222 | 5343 | 1880 |
| 48 | ATAC | all | down | 1000 | hmg4_vs_spt16 | 9. Transcription 3' 3 | 5.1E-06 | 1.6E-05 | 1.98 | 4.16 | 19 | 457 | 1.09 | 58 | 5343 | 2675 |
| 49 | ATAC | all | up | 200 | hmg4_vs_spt16 | 14. Low signal 1 | 2.5E-06 | 0.00002 | 2.01 | 15.45 | 19 | 123 | 4.35 | 247 | 5677 | 1880 |
| 50 | ATAC | all | down | 200 | hmg4_vs_spt16 | 7. Transcription 3' 1 | 2.5E-06 | 2.4E-05 | 2.8 | 13.41 | 11 | 82 | 2.17 | 124 | 5718 | 3424 |
| 51 | ATAC | all | down | 200 | hmg4_vs_spt16 | 2. Enhancer 1 | 3E-06 | 2.4E-05 | -2.37 | 7.32 | 6 | 82 | 29.03 | 1660 | 5718 | 2785 |

q

| | A | B | C | D | E | F | G | H | I | J | K | L | M | N | O | P | Q | R |
|---|---|---|---|---|---|---|---|---|---|---|---|---|---|---|---|---|---|---|
| 1 | GE | ET | PF | FC | TV | COMP | tgt | pval | padj | L2OR | pt_da | ov_da | tot_da | pt_nda | ov_nda | tot_nda | tgt_id | genes_id |
| 2 | BP | mRNA | Null | up | 1000 | hmg4_vs_ctl | carboxylic acid metabolic process | 4.1E-30 | 2E-27 | 2.54 | 20.55 | 82 | 399 | 4.26 | 407 | 9553 | GO:0019752 | WBGene00019322/WBGer |
| 3 | BP | mRNA | Null | up | 1000 | hmg4_vs_ctl | oxoacid metabolic process | 4.7E-30 | 2E-27 | 2.54 | 20.55 | 82 | 399 | 4.27 | 408 | 9553 | GO:0043436 | WBGene00019322/WBGer |
| 4 | BP | mRNA | Null | up | 1000 | hmg4_vs_spt16 | carboxylic acid metabolic process | 7.9E-23 | 2.6E-20 | 2.33 | 18.25 | 69 | 378 | 4.26 | 407 | 9553 | GO:0019752 | WBGene00019322/WBGer |
| 5 | BP | mRNA | Null | up | 1000 | hmg4_vs_spt16 | oxoacid metabolic process | 9E-23 | 2.6E-20 | 2.32 | 18.25 | 69 | 378 | 4.27 | 408 | 9553 | GO:0043436 | WBGene00019322/WBGer |
| 6 | BP | mRNA | Null | up | 1000 | hmg4_vs_spt16 | organic acid metabolic process | 1E-22 | 2.6E-20 | 2.26 | 19.05 | 72 | 378 | 4.67 | 446 | 9553 | GO:0006082 | WBGene00019322/WBGer |
| 7 | BP | mRNA | Null | down | 200 | hmg4_vs_ctl | collagen and cuticulin-based cuticle develc | 1.1E-18 | 9.4E-17 | 6.2 | 25 | 13 | 52 | 0.45 | 43 | 9553 | GO:0040002 | WBGene00001066/WBGer |
| 8 | BP | mRNA | Null | down | 200 | hmg4_vs_ctl | molting cycle, collagen and cuticulin-based | 7.8E-19 | 9.4E-17 | 5.35 | 30.77 | 16 | 52 | 1.08 | 103 | 9553 | GO:0018996 | WBGene00001066/WBGer |
| 9 | BP | mRNA | Null | down | 200 | hmg4_vs_ctl | molting cycle | 1.2E-18 | 9.4E-17 | 5.3 | 30.77 | 16 | 52 | 1.11 | 106 | 9553 | GO:0042303 | WBGene00001066/WBGer |
| 10 | BP | mRNA | Null | down | 200 | spt16_vs_ctl | cuticle development | 2.7E-18 | 6.9E-16 | 5.73 | 21.21 | 14 | 66 | 0.5 | 48 | 9553 | GO:0042335 | WBGene00005018/WBGer |
| 11 | BP | mRNA | Null | down | 200 | spt16_vs_ctl | molting cycle, collagen and cuticulin-based | 5.5E-17 | 7E-15 | 4.87 | 24.24 | 16 | 66 | 1.08 | 103 | 9553 | GO:0018996 | WBGene00005018/WBGer |
| 12 | BP | mRNA | Null | down | 200 | spt16_vs_ctl | molting cycle | 8.2E-17 | 7.1E-15 | 4.83 | 24.24 | 16 | 66 | 1.11 | 106 | 9553 | GO:0042303 | WBGene00005018/WBGer |
| 13 | BP | mRNA | Null | down | 200 | spt16_vs_ctl | cuticle development involved in collagen a | 6.9E-16 | 4.5E-14 | 6.13 | 16.67 | 11 | 66 | 0.28 | 27 | 9553 | GO:0042338 | WBGene00005018/WBGer |
| 14 | BP | mRNA | Null | down | 200 | hmg4_vs_ctl | cuticle development involved in collagen a | 2.7E-15 | 1.6E-13 | 6.38 | 19.23 | 10 | 52 | 0.28 | 27 | 9553 | GO:0042338 | WBGene00001066/WBGer |
| 15 | BP | mRNA | Null | down | 1000 | spt16_vs_ctl | collagen and cuticulin-based cuticle develc | 6.7E-16 | 2.6E-13 | 3.79 | 5.9 | 22 | 373 | 0.45 | 43 | 9553 | GO:0040002 | WBGene00005018/WBGer |

r

| | A | B | C | D | E | F | G | H | I | J | K | L | M | N | O | P | Q | R |
|---|---|---|---|---|---|---|---|---|---|---|---|---|---|---|---|---|---|---|
| 1 | GE | ET | PF | FC | TV | COMP | tgt | pval | padj | L2OR | pt_da | ov_da | tot_da | pt_nda | ov_nda | tot_nda | tgt_id | genes_id |
| 2 | KEGG | mRNA | Null | up | 1000 | hmg4_vs_ctl | Carbon metabolism | 8.3E-14 | 7.5E-12 | 2.41 | 17.17 | 40 | 233 | 3.76 | 108 | 2875 | cel01200 | WBGene00014095/WBGe |
| 3 | KEGG | ATAC | all | down | 1000 | spt16_vs_ctl | Aminoacyl-tRNA biosynthesis | 9E-11 | 8.8E-09 | -4.91 | 0.49 | 1 | 203 | 12.94 | 372 | 2875 | cel00970 | WBGene00013447 |
| 4 | KEGG | mRNA | Null | up | 1000 | hmg4_vs_ctl | Pyruvate metabolism | 1.7E-09 | 5.1E-08 | 2.93 | 8.15 | 19 | 233 | 1.15 | 33 | 2875 | cel00620 | WBGene00000115/WBGe |
| 5 | KEGG | mRNA | Null | up | 1000 | hmg4_vs_ctl | Valine, leucine and isoleucine degradatior | 1.6E-09 | 5.1E-08 | 2.57 | 9.87 | 23 | 233 | 1.81 | 52 | 2875 | cel00280 | WBGene00000115/WBGe |
| 6 | KEGG | mRNA | Null | up | 1000 | hmg4_vs_ctl | Aminoacyl-tRNA biosynthesis | 3.5E-09 | 8E-08 | -3.09 | 1.72 | 4 | 233 | 12.94 | 372 | 2875 | cel00970 | WBGene00006617/WBGe |
| 7 | KEGG | mRNA | Null | up | 1000 | hmg4_vs_ctl | Glycolysis / Gluconeogenesis | 7.6E-09 | 1.4E-07 | 2.66 | 8.58 | 20 | 233 | 1.46 | 42 | 2875 | cel00010 | WBGene00000115/WBGe |
| 8 | KEGG | ATAC | all | down | 1000 | hmg4_vs_ctl | Aminoacyl-tRNA biosynthesis | 7.3E-09 | 6.4E-07 | -4.62 | 0.6 | 1 | 167 | 12.94 | 372 | 2875 | cel00970 | WBGene00013447 |
| 9 | KEGG | mRNA | Null | up | 200 | hmg4_vs_ctl | Carbon metabolism | 1.5E-08 | 6.5E-07 | 3.81 | 35.48 | 11 | 31 | 3.76 | 108 | 2875 | cel01200 | WBGene00014095/WBGe |
| 10 | KEGG | mRNA | Null | up | 1000 | hmg4_vs_spt16 | Aminoacyl-tRNA biosynthesis | 1.1E-08 | 1E-06 | -2.79 | 2.11 | 5 | 237 | 12.94 | 372 | 2875 | cel00970 | WBGene00004189/WBGe |
| 11 | KEGG | mRNA | Null | down | 1000 | spt16_vs_ctl | Protein processing in endoplasmic reticulu | 1.9E-08 | 1.6E-06 | 1.9 | 15.76 | 32 | 203 | 4.77 | 137 | 2875 | cel04141 | WBGene00003963/WBGe |
| 12 | KEGG | mRNA | Null | up | 1000 | spt16_vs_ctl | Carbon metabolism | 3E-08 | 2.7E-06 | 2.1 | 14.36 | 26 | 181 | 3.76 | 108 | 2875 | cel01200 | WBGene00014095/WBGe |
| 13 | KEGG | mRNA | Null | up | 1000 | hmg4_vs_spt16 | Carbon metabolism | 6E-08 | 2.8E-06 | 1.89 | 12.66 | 30 | 237 | 3.76 | 108 | 2875 | cel01200 | WBGene00003214/WBGe |
| 14 | KEGG | mRNA | Null | up | 1000 | hmg4_vs_ctl | Citrate cycle (TCA cycle) | 2.1E-07 | 3.2E-06 | 2.67 | 6.87 | 16 | 233 | 1.15 | 33 | 2875 | cel00020 | WBGene00011232/WBGe |
| 15 | KEGG | mRNA | Null | up | 1000 | spt16_vs_ctl | Glycolysis / Gluconeogenesis | 1.1E-07 | 4.7E-06 | 2.71 | 8.84 | 16 | 181 | 1.46 | 42 | 2875 | cel00010 | WBGene00010790/WBGe |

s

| | A | B | C | D | E | F | G | H | I | J | K | L | M | N | O | P |
|---|---|---|---|---|---|---|---|---|---|---|---|---|---|---|---|---|
| 1 | ET | PF | FC | TV | COMP | tgt | pval | padj | L2OR | pt_da | ov_da | tot_da | pt_nda | ov_nda | tot_nda | tot_tgt |
| 100 | mRNA | Null | down | 200 | hmg4_vs_ctl | mRNA__Null__down__200__spt16_vs_ctl | 3.2E-81 | 1.2E-79 | 6.57 | 73.45 | 83 | 113 | 2.8 | 43 | 1533 | 126 |
| 101 | ATAC | all | down | 200 | hmg4_vs_ctl | ATAC__prom__down__1000__hmg4_vs_ctl | 1.2E-78 | 1.8E-77 | 4.63 | 54.5 | 103 | 189 | 4.61 | 293 | 6362 | 396 |
| 102 | ATAC | prom | down | 1000 | hmg4_vs_ctl | ATAC__all__down__200__hmg4_vs_ctl | 1.3E-78 | 2.4E-77 | 4.63 | 26.01 | 103 | 396 | 1.4 | 86 | 6151 | 189 |
| 103 | ATAC | prom | down | 1000 | hmg4_vs_ctl | ATAC__prom__down__1000__spt16_vs_ctl | 8.6E-77 | 1.2E-75 | 3.5 | 37.88 | 150 | 396 | 5.12 | 315 | 6151 | 468 |
| 104 | ATAC | all | down | 1000 | spt16_vs_ctl | ATAC__all__down__1000__hmg4_vs_ctl | 5.5E-76 | 6.6E-75 | 2.45 | 35.89 | 290 | 808 | 9.28 | 470 | 5062 | 763 |
| 105 | mRNA | Null | down | 1000 | spt16_vs_ctl | mRNA__Null__down__1000__hmg4_vs_ctl | 4.9E-74 | 1.2E-72 | 3.13 | 56.92 | 288 | 506 | 13.12 | 155 | 1181 | 474 |
| 106 | ATAC | prom | down | 1000 | spt16_vs_ctl | ATAC__prom__down__1000__hmg4_vs_ctl | 1.2E-73 | 1.7E-72 | 3.41 | 32.05 | 150 | 468 | 4.24 | 243 | 5735 | 396 |
| 107 | ATAC | all | down | 200 | spt16_vs_ctl | ATAC__distNC__down__200__spt16_vs_ctl | 1.2E-72 | 1.7E-71 | 1E+99 | 25 | 45 | 180 | 0 | 0 | 6091 | 45 |
| 108 | ATAC | distNC | down | 200 | spt16_vs_ctl | ATAC__all__down__200__spt16_vs_ctl | 7E-73 | 2.5E-71 | 1E+99 | 100 | 45 | 45 | 2.14 | 135 | 6295 | 180 |
| 109 | mRNA | Null | down | 1000 | spt16_vs_ctl | mRNA__Null__down__200__spt16_vs_ctl | 6.1E-72 | 1.1E-70 | 1E+99 | 24.9 | 126 | 506 | 0 | 0 | 1181 | 126 |
| 110 | mRNA | Null | down | 200 | spt16_vs_ctl | mRNA__Null__down__1000__spt16_vs_ctl | 6.1E-72 | 1.5E-70 | 1E+99 | 100 | 126 | 126 | 24.34 | 380 | 1561 | 506 |
| 111 | ATAC | prom | up | 200 | hmg4_vs_spt16 | ATAC__all__up__200__hmg4_vs_spt16 | 2.2E-71 | 7.9E-70 | 1E+99 | 100 | 40 | 40 | 1.37 | 79 | 5752 | 119 |
| 112 | ATAC | all | up | 200 | hmg4_vs_spt16 | ATAC__prom__up__200__hmg4_vs_spt16 | 3.3E-71 | 8E-70 | 1E+99 | 33.61 | 40 | 119 | 0 | 0 | 5612 | 40 |
| 113 | mRNA | Null | up | 1000 | spt16_vs_ctl | mRNA__Null__down__1000__hmg4_vs_spt16 | 5.9E-71 | 1.4E-69 | 3 | 59.72 | 295 | 494 | 15.59 | 186 | 1193 | 514 |
| 114 | ATAC | distNC | down | 200 | spt16_vs_ctl | ATAC__distNC__down__1000__spt16_vs_ctl | 3E-68 | 7.2E-67 | 1E+99 | 100 | 45 | 45 | 2.81 | 177 | 6295 | 222 |

t

| | A | B | C | D | E | F | G | H | I | J | K | L | M | N | O | P |
|---|---|---|---|---|---|---|---|---|---|---|---|---|---|---|---|---|
| 1 | ET | PF | FC | TV | COMP | tgt | pval | padj | L2OR | pt_da | ov_da | tot_da | pt_nda | ov_nda | tot_nda | consensus |
| 29 | mRNA | Null | down | 1000 | hmg4_vs_ctl | zip-8 | 6.3E-07 | 5.9E-05 | 0.9 | 19.2 | 91 | 474 | 11.31 | 127.3 | 1126 | ATTACGTAAT |
| 30 | mRNA | Null | up | 1000 | hmg4_vs_spt16 | mls-2 | 1.5E-06 | 0.00011 | -0.65 | 59.47 | 289 | 486 | 69.79 | 736.3 | 1055 | BKTTAATTGBT |
| 31 | mRNA | Null | up | 200 | spt16_vs_ctl | sex-1 | 2.4E-06 | 0.00091 | 1.65 | 49.32 | 36 | 73 | 23.66 | 364.2 | 1539 | RAGGTCAM |
| 32 | mRNA | Null | down | 1000 | spt16_vs_ctl | mls-2 | 8.4E-06 | 0.001 | -0.6 | 60.08 | 304 | 506 | 69.46 | 793.9 | 1143 | BKTTAATTGBT |
| 33 | mRNA | Null | up | 1000 | hmg4_vs_ctl | pag-3 | 5.7E-05 | 0.0021 | -2.37 | 98.1 | 516 | 526 | 99.61 | 1066.8 | 1071 | NAAATCWSNG |
| 34 | mRNA | Null | down | 1000 | hmg4_vs_ctl | ast-1 | 2.9E-05 | 0.0022 | -0.71 | 78.9 | 374 | 474 | 85.94 | 967.7 | 1126 | NACCGGAARY |
| 35 | mRNA | Null | down | 200 | hmg4_vs_ctl | ceh-38 | 0.00002 | 0.0037 | -1.33 | 21.24 | 24 | 113 | 40.49 | 603.3 | 1490 | WNCGATCGATAAHN |
| 36 | mRNA | Null | down | 1000 | spt16_vs_ctl | lin-1 | 6.3E-05 | 0.0058 | -0.53 | 59.29 | 300 | 506 | 67.75 | 774.4 | 1143 | NRCCGGAART |
| 37 | mRNA | Null | down | 1000 | hmg4_vs_spt16 | jun-1 | 3.4E-05 | 0.0064 | 0.57 | 35.28 | 181 | 513 | 26.93 | 278.2 | 1033 | ATGACTCAT |
| 38 | mRNA | Null | down | 1000 | spt16_vs_ctl | nhr-2 | 0.00012 | 0.0074 | 0.98 | 8.3 | 42 | 506 | 4.4 | 50.3 | 1143 | ATGTAGGTCAMN |
| 39 | mRNA | Null | down | 1000 | spt16_vs_ctl | nhr-85 | 0.00012 | 0.0074 | 0.98 | 8.3 | 42 | 506 | 4.4 | 50.3 | 1143 | ATGTAGGTCAMN |
| 40 | mRNA | Null | up | 1000 | hmg4_vs_spt16 | ref-1 | 0.00012 | 0.0074 | 0.61 | 23.46 | 114 | 486 | 16.67 | 175.9 | 1055 | ACACGTGC |
| 41 | mRNA | Null | down | 200 | hmg4_vs_spt16 | ztf-14 | 2.1E-05 | 0.008 | 1.35 | 46.59 | 41 | 88 | 25.47 | 369.8 | 1452 | VGACCACCCA |
| 42 | ATAC | all | down | 1000 | hmg4_vs_spt16 | ztf-16 | 2.3E-05 | 0.0087 | 0.69 | 24.29 | 111 | 457 | 16.58 | 840.3 | 5068 | NYYCHGGATN |
| 43 | mRNA | Null | up | 1000 | hmg4_vs_spt16 | pal-1 | 0.00018 | 0.0095 | -0.53 | 66.67 | 324 | 486 | 74.31 | 784 | 1055 | GTAATWAA |
| 44 | mRNA | Null | up | 1000 | hmg4_vs_ctl | efl-3 | 0.00029 | 0.0097 | 1.64 | 2.85 | 15 | 526 | 0.98 | 10.5 | 1071 | TTTGGCGGGAAA |
| 45 | mRNA | Null | up | 1000 | hmg4_vs_spt16 | fos-1 | 0.00026 | 0.011 | -0.68 | 13.37 | 65 | 486 | 19.81 | 209 | 1055 | TGACTCA |

u

| | A | B | C | D | E | F | G | H | I | J | K | L | M | N | O | P |
|---|---|---|---|---|---|---|---|---|---|---|---|---|---|---|---|---|
| 1 | ET | PF | FC | TV | COMP | tgt | pval | padj | L2OR | pt_da | ov_da | tot_da | pt_nda | ov_nda | tot_nda | tot_tgt |
| 100 | both_ATAC | all | down | 1000 | spt16_vs_ctl | both_ATAC__all__down__1000__spt16_vs_ctl | 2.1E-82 | 1.7E-80 | 1E+99 | 100 | 30 | 30 | 0 | 0 | 6346 | 30 |
| 101 | both_ATAC | all | down | 1000 | hmg4_vs_spt16 | both_ATAC__all__down__1000__hmg4_vs_spt16 | 3.6E-81 | 2.9E-79 | 1E+99 | 100 | 30 | 30 | 0 | 0 | 5770 | 30 |
| 102 | mRNA | Null | down | 200 | hmg4_vs_ctl | mRNA__Null__down__200__spt16_vs_ctl | 9.5E-80 | 3.9E-78 | 6.46 | 73.45 | 83 | 113 | 3.01 | 46 | 1527 | 126 |
| 103 | ATAC | all | down | 200 | hmg4_vs_ctl | ATAC__prom__down__1000__hmg4_vs_ctl | 1.1E-77 | 1.9E-76 | 4.6 | 53.68 | 102 | 190 | 4.54 | 295 | 6494 | 397 |
| 104 | ATAC | prom | down | 1000 | hmg4_vs_ctl | ATAC__all__down__200__hmg4_vs_ctl | 1.1E-77 | 2.4E-76 | 4.6 | 25.69 | 102 | 397 | 1.4 | 88 | 6287 | 190 |
| 105 | ATAC | prom | down | 1000 | hmg4_vs_ctl | ATAC__prom__down__1000__spt16_vs_ctl | 3.8E-75 | 6.2E-74 | 3.46 | 37.28 | 148 | 397 | 5.11 | 321 | 6287 | 469 |
| 106 | ATAC | all | down | 200 | spt16_vs_ctl | ATAC__distNC__down__200__spt16_vs_ctl | 1.6E-74 | 2.6E-73 | 1E+99 | 25.41 | 46 | 181 | 0 | 0 | 6195 | 46 |
| 107 | ATAC | distNC | down | 200 | spt16_vs_ctl | ATAC__all__down__200__spt16_vs_ctl | 1.6E-74 | 6.4E-73 | 1E+99 | 100 | 46 | 46 | 2.13 | 135 | 6330 | 181 |
| 108 | both_ATAC | all | up | 1000 | hmg4_vs_spt16 | both_ATAC__all__up__1000__hmg4_vs_spt16 | 2.8E-74 | 2.3E-72 | 1E+99 | 100 | 27 | 27 | 0 | 0 | 5773 | 27 |
| 109 | ATAC | prom | down | 1000 | spt16_vs_ctl | ATAC__prom__down__1000__hmg4_vs_ctl | 1.5E-72 | 2.5E-71 | 3.39 | 31.56 | 148 | 469 | 4.2 | 248 | 5907 | 397 |
| 110 | mRNA | Null | down | 1000 | spt16_vs_ctl | mRNA__Null__down__200__spt16_vs_ctl | 5.9E-72 | 1.6E-70 | 1E+99 | 25.3 | 128 | 506 | 0 | 0 | 1146 | 126 |
| 111 | ATAC | all | up | 200 | hmg4_vs_spt16 | ATAC__prom__up__200__hmg4_vs_spt16 | 1E-70 | 2.8E-69 | 1E+99 | 32.52 | 40 | 123 | 0 | 0 | 5677 | 40 |
| 112 | ATAC | prom | up | 200 | hmg4_vs_spt16 | ATAC__all__up__200__hmg4_vs_spt16 | 1E-70 | 4.2E-69 | 1E+99 | 100 | 40 | 40 | 1.44 | 83 | 5760 | 123 |
| 113 | mRNA | Null | up | 1000 | spt16_vs_ctl | mRNA__Null__down__1000__hmg4_vs_spt16 | 3.8E-70 | 1E-68 | 2.97 | 61.74 | 305 | 494 | 17.06 | 197 | 1155 | 514 |
| 114 | ATAC | distNC | down | 1000 | spt16_vs_ctl | ATAC__distNC__down__200__spt16_vs_ctl | 9.7E-69 | 2.6E-67 | 1E+99 | 19.66 | 46 | 234 | 0 | 0 | 6142 | 46 |
